# DNA Nicks Drive Massive Expansions of (GAA)_n_ Repeats

**DOI:** 10.1101/2024.06.12.598717

**Authors:** Liangzi Li, W. Shem Scott, Alexandra N. Khristich, Jillian F. Armenia, Sergei M. Mirkin

## Abstract

Over 50 hereditary degenerative disorders are caused by expansions of short tandem DNA repeats (STRs). (GAA)_n_ repeat expansions are responsible for Friedreich’s ataxia as well as late-onset cerebellar ataxias (LOCAs). Thus, the mechanisms of (GAA)_n_ repeat expansions attract broad scientific attention. To investigate the role of DNA nicks in this process, we utilized a CRISPR-Cas9 nickase system to introduce targeted nicks adjacent to the (GAA)_n_ repeat tract. We found that DNA nicks 5’ of the (GAA)_100_ run led to a dramatic increase in both the rate and scale of its expansion in dividing cells. Strikingly, they also promoted large-scale expansions of carrier- and large normal-size (GAA)_n_ repeats, recreating, for the first time in a model system, the expansion events that occur in human pedigrees. DNA nicks 3’ of the (GAA)_100_ repeat led to a smaller but significant increase in the expansion rate as well. Our genetic analysis implies that in dividing cells, conversion of nicks into double-strand breaks (DSBs) during DNA replication followed by DSB or fork repair leads to repeat expansions. Finally, we showed that 5’ strand nicks increase expansion frequency in non-dividing yeast cells, albeit to a lesser extent that in dividing cells.

## Introduction

Up to 6% of the human genome is comprised of or derived from STRs^1,2^, which are 2-9 bp long repetitive DNA sequences. STRs exhibit substantial length-polymorphism, which is believed to play a role in gene regulation^3^. Except for structural chromosomal elements such as telomeric repeats^4^, STRs normally consist of a limited number of repeats rarely exceeding 30 copies.

Occasionally, however, a genic STR can expand far beyond its normal length ultimately leading to disease. Over 50 hereditary human disorders, cumulatively called repeat expansion diseases (REDs), are linked to such unruly STR expansions^5,6^. This number continues to grow owing to advances in long-read sequencing^7^. Recent additions include expansions of (AAGGG)_n_ repeats in the *RFC1* gene causing cerebellar ataxia with neuropathy and vestibular areflexia syndrome (CANVAS)^8^, expansions of (GAA)_n_ repeats in the *FGF14* gene responsible for LOCA^9^, and expansions of (GGC)_n_ repeats in the *ZFHX3* gene linked to spinocerebellar ataxia type 4 (SCA4)^10^.

Since the characterization of the first RED – fragile X syndrome (FXS) – in 1991^11–13^, an STR progression from normal to disease length in human pedigrees was extensively characterized^14–19^. It starts from the so-called long-normal alleles that carry 20-to-40 repeats with several interruptions. Subsequent loss of those interruptions brings in carrier alleles containing 25-to-35 non-interrupted repetitive runs. Finally, large-scale repeat expansions in carrier alleles give rise to pathogenic alleles that contain hundreds or even thousands of extra repeats. During intergenerational transmissions, these expansion events occur in rapidly dividing pre-meiotic germ cells or at the early steps of embryonic development (reviewed in ref. 20,21). An inherited pathogenic allele often undergoes further repeat expansions in patients’ somatic cells, including post-mitotic cells^22–25^. It is believed that sequential accumulation of extra repeats in somatic cells accounts for the late onset of most REDs^23,26–29^.

Genetic control of repeat expansions was studied in human pedigrees as well as in model systems. Several genome-wide association studies (GWAS) identified DNA repair genes, including mismatch repair genes, as modifiers of disease progression^30–32^. More recently, *FAN1* gene, encoding a 5’-flap endonuclease, emerged as the strongest modifier of REDs^33,34^.

Molecular genetic analysis conducted in model systems, including bacteria, yeast, Drosophila and mice implicated DNA replication^35–38^, DNA repair^39–41^ and DNA recombination^42–45^ (reviewed in ref. 46) in the expansion process. It has also become clear that genetic pathways of repeat expansions are not mutually exclusive but can contribute to the process at different developmental stages and tissues.

One common denominator for all these mechanisms could be single-strand breaks (SSBs) or DNA nicks. DNA nicks are among the most common DNA lesions in metabolically active cells^47,48^. They can arise directly from DNA damaging agents, such as reactive oxygen species. Repair of DNA nicks involves the formation of 5’-flaps that are then removed by flap endonucleases, such as FAN1. At the same time, DNA nicks play essential roles in normal genome functioning. For example, their presence in the nascent lagging strand during DNA replication^49,50^ allows mismatch repair (MMR) machinery to discriminate between the template and nascent DNA strands^51^. They can also initiate homologous recombination (HR)^52^ (reviewed in ref. 47).

Studying the mouse model for Huntington’s disease, the McMurray lab showed that somatic repeat expansions occur during removal of 8-oxoguanine by the base excision repair enzyme 7,8-dihydro-8-oxoguanine-DNA glycosylase (OGG1)^53^. Oxidative stress stimulated somatic repeat expansions^54^, while synthetic antioxidant XJB-5-131 suppressed somatic expansions^27,55^. Altogether this led them to propose the “toxic oxidation cycle” model, stipulating that somatic repeat expansions occur during base excision repair (BER) of oxidative DNA lesions. Removal of oxidated bases by concerted actions of DNA glycosylases and AP endonuclease creates DNA nicks. In the context of a structure-prone repeat, repair synthesis by DNA polymerase could lead to the formation of an alternative DNA structure in the displaced flap, ultimately producing an expanded repeat, which then accumulates even more oxidative damage fueling this toxic cycle^56–60^. Supporting this idea, strand displacement activity was shown to enable DNA polymerase beta to expand CAG/CTG triplet repeats at DNA nicks *in vitro*^61,62^. This consideration applies to other expandable repeats as well. For example, BER of alkylated DNA damage was shown to destabilize expanded Friedreich’s ataxia GAA repeats^63^, while omaveloxolone – an activator of nuclear factor erythroid 2-related factor 2 counteracting oxidative damage – became the first FDA-approved drug counteracting repeat expansion in Friedreich’s ataxia patients^64^. We have shown that GAA repeat expansions in quiescent yeast cells was exacerbated by the loss of Exo1 involved in DNA end resection, pointing to the role DNA nick repair^39^.

Developments in genome editing techniques allow one to generate nicks *in vivo* at precise genomic positions^65,66^. A pioneering study of the Dion lab demonstrated that induction of multiple DNA nicks within the (CAG)_n_ repeat in a reporter human cassette by the CRISPR-Cas9 nickase promoted its contractions^67^. More recently, induction of DNA nicks adjacent to the (G_4_C_2_)_n_ repeat in the Amyotrophic Lateral Sclerosis (ALS) and Frontotemporal Dementia (FTD) mice triggered both its expansions and contractions^68^. These data show destabilization of expandable repeats by targeted DNA nicks, which has broad biomedical potential. The molecular mechanisms and genetic controls of the destabilization remain to be understood.

To address these mechanisms, we investigated the effects of DNA nicks generated by CRISPR-Cas9 nickases on GAA repeat expansions in our yeast intronic system. We found that nicks 5’ to the repeat led to extremely frequent and very large-scale expansions of the pathogenic (GAA)_100_ repeat, as well as the premutation (GAA)_40_ and long-normal (GAA)_33_ repeats in dividing cells. Furthermore, we observed a substantial increase in the frequency of repeat expansions in quiescent yeast cells. Extensive candidate gene analysis revealed that nick-mediated repeat expansions in dividing cells were steered by either homologous recombination or replication fork repair pathways, depending on the position of a nick.

## Results

### Nicks adjacent to the GAA repeat induce large-scale expansions in dividing yeast cells

To study the effects of DNA nicks on repeat expansions, we utilized our intronic system for GAA repeat expansions in yeast^37,38^. In brief, the (GAA)_100_ repeat was inserted into an artificial intron of the *URA3* gene; the total length of this intron was 974 bp. The cassette containing split *UR*-*(GAA)_100_*-*A3* gene followed by the *TRP1* gene was integrated into chromosome III, roughly 1 kb downstream of ARS306 replication origin (Figure 1A). Since yeast cannot splice introns over ∼1 kb, repeat expansions of more than ∼20 copies inactivate the *URA3* gene, making cells resistant to 5-fluoroorotic acid (5-FOA) (Figures 1A&S1A). To induce targeted nicks, we created a CRISPR-Cas9 dual plasmid system inspired by Nishida *et al.*^66^ One of these plasmids expressed either nCas9-D10A or nCas9-H840A nickase under the inducible *GAL1* promoter, while the other plasmid expressed the gRNA under the control of *SNR52* RNA polymerase III (RNAPIII) promoter (Figure S1B). The combination of a nCas9 plasmid and a gRNA plasmid was co-transformed into yeast strains carrying the *UR-(GAA)_100_-A3* cassette. We identified two protospacer adjacent motif (PAM) sites on either side of the (GAA)_100_ repeat tract. Thus, between two nickases and two PAM sites, we were able to induce four types of nicks around the repeat: 5’ GAA-strand and 5’ TTC-strand nicks were induced by nCas9-D10A, while 3’ GAA-strand and 3’ TTC-strand nicks were induced by nCas9-H840A (Figure 1B). Nicks were located 6-to-7 nucleotides from the repeat tract (Figure S1C).

**Figure 1.**
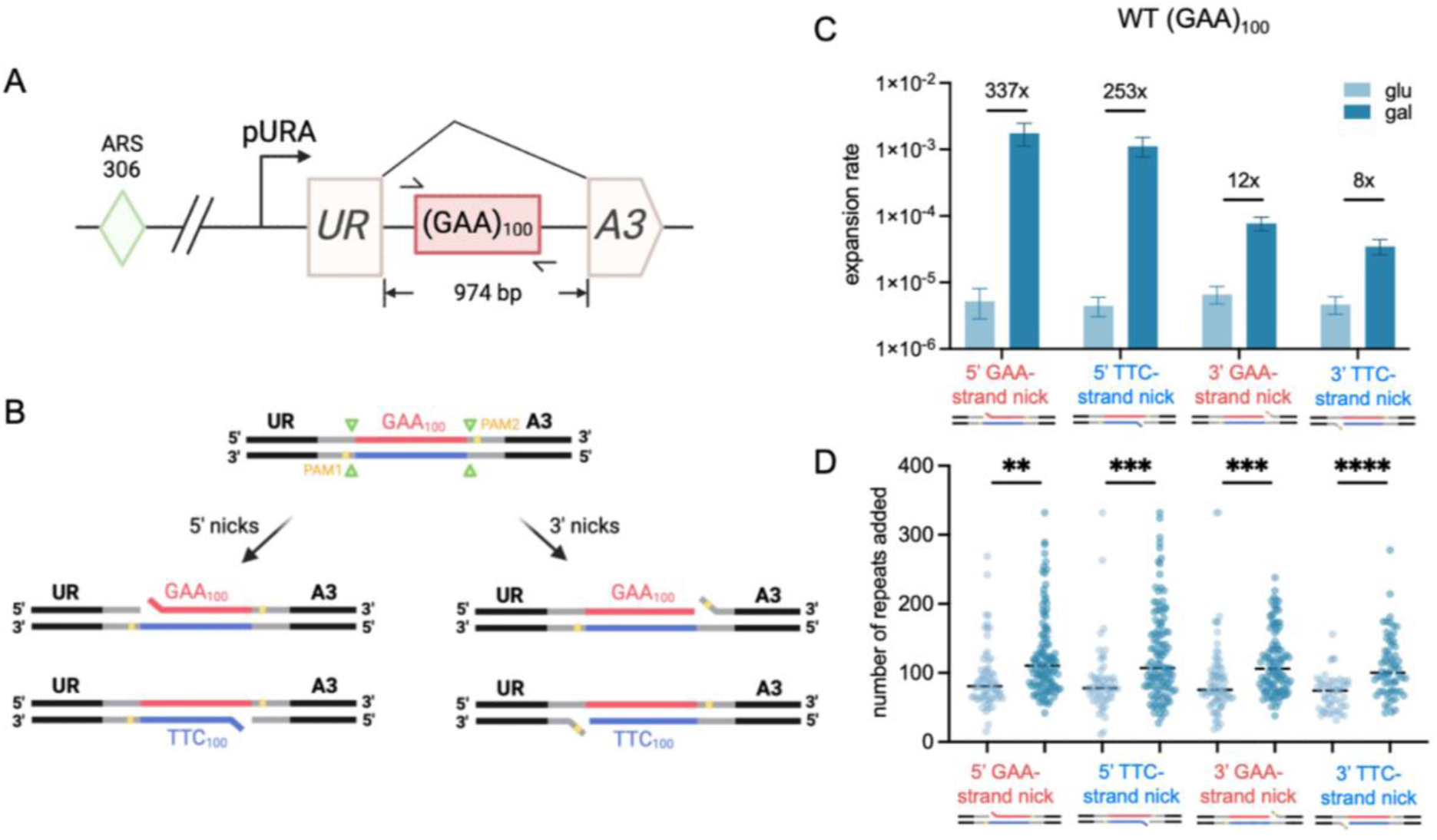
An experimental system to study nick-mediated repeat expansion in yeast. (A) The integrated *URA3* reporter cassette for selection of large-scale (GAA)_100_ repeat expansions. The (GAA)_100_ repeat located within an artificial intron of the *URA3* gene which has the total length of 974 bp. A TRP1 gene is located downstream as a selectable marker for cassette integration. The entire cassette is integrated ∼1kb downstream of an early-firing origin of replication ARS306 on chromosome III. (B) Schematic of nCas9 cut sites adjacent to the (GAA)_100_ repeat. Two PAM sites, one on each end of the repeat, were identified to allow for nCas9 nicking. Four types of nicks were induced, which were categorized into 5’ GAA or TTC-strand nicks or 3’ GAA or TTC-strand nicks. (C) Expansion rate of the WT strains carrying all four combinations of nickase plasmids without (glu, light blue) or with (gal, dark blue) galactose induction of Cas9 expression. Numbers on top of each genotype show fold change of expansion rate between glucose and galactose group. Error bars represent 95% confidence intervals. (D) Expansion distribution of WT strains carrying all four combinations of nickase plasmids with (dark blue) or without (light blue) galactose induction of Cas9 expression. Dashed lines represent median of distribution. Statistical significance between glucose and galactose groups was calculated via Kolmogorov–Smirnov test.

To assess the effect of nicks on (GAA)_100_ repeat expansion, cells with the nickase plasmids were cultured on the media containing glucose or galactose as the carbon source. Individual colonies grown on glucose or galactose media were picked and after serial dilutions plated on both non-selective (YPD) and selective (5-FOA) media followed by growth at 30 °C for 3 days. Repeat expansions in 5-FOA resistant (5-FOA^r^) colonies were confirmed by PCR. Expansion rates were calculated using *FluCalc*^69^. The distribution of expansion lengths was also measured, and statistical significance of differences between individual distributions was evaluated via Kolmogorov–Smirnov test (KS-test).

Figure 1C shows that expansion rate of the (GAA)_100_ repeat increased significantly when any of the four nicks was targeted to the repeat. Nicks 5’ to the repeat on either strand dramatically increased its expansion rate by over two orders of magnitude, while nicks 3’ of the repeat increased expansion rate by roughly one order of magnitude. 5’ nicks led to the biggest increases in the expansion rate we have ever observed in our yeast system. In addition to the rates, the scale of repeat expansions also significantly increased by all four types of nicks (Figure 1D). Taken together, we showed that nicks adjacent to the (GAA)_100_ repeat dramatically increased both expansion rate and length.

### 5’ GAA-strand nicks induce expansions of premutation- and long-normal-size repeats

For Friedreich’s ataxia, most unaffected individuals have less than 10 GAA repeats in the first intron of their *FXN* gene^70^, premutation alleles contain 34-to-66 repeats, while alleles longer than 66 units are considered to be pathogenic (Figure 2A)^19,71–73^. In a few cases, hyper-expansions were observed in individuals carrying 34 GAA repeats^71^. In our previous studies, we were unable to observe large-scale expansions for non-pathogenic repeats, even though the sensitivity of our system allows for the detection of events occurring at a rate of 10^-7^ per cell per generation^38^. The extremely high increase in expansion rates upon nick induction led us to revisit the possibility of large-scale expansions for premutation and long-normal repeat alleles.

**Figure 2.**
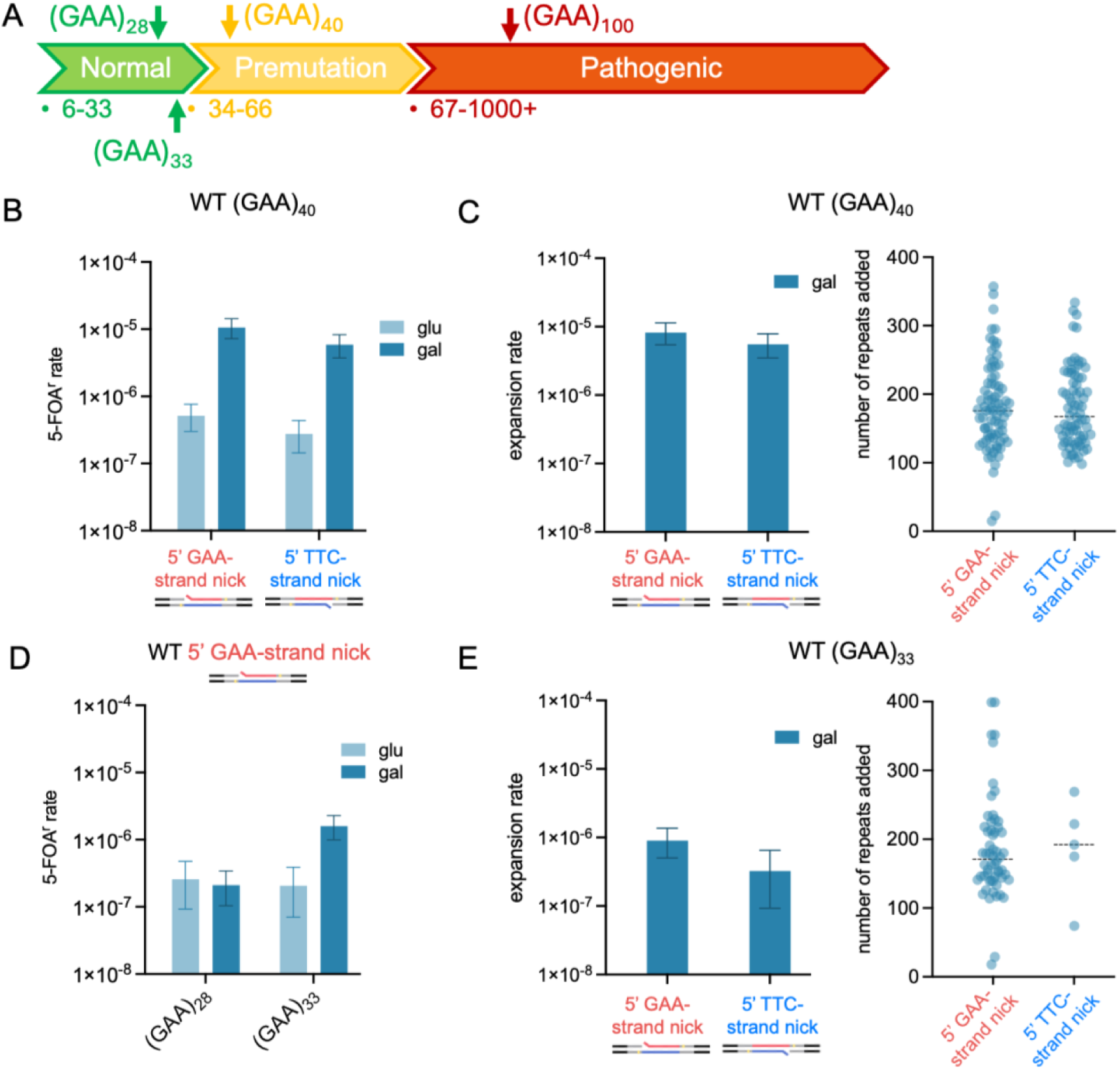
5’ nick-mediated expansion in premutation and long-normal (GAA)_n_ alleles. (A) Thresholds of (GAA)_n_ repeat associated with Friedreich’s ataxia. Normal alleles range from 6 to 33 repeats, premutation alleles range between 34 to 66 repeats and pathogenic alleles are larger than 67 repeats. (B) Effect of 5’ nicks on 5-FOA^r^ rate of WT (GAA)_40_ strains without (glu, light blue) or with (gal, dark blue) galactose induction of nCas9-D10A expression. Error bars represent 95% confidence intervals. (C) Expansion rate (left) and scale (right) of WT (GAA)_40_ strains with galactose induction of nCas9-D10A expression. Left: Error bars represent 95% confidence intervals. Right: Dashed lines represent median of distribution. (D) Effect of 5’ GAA-strand nick on 5-FOA^r^ rate of WT (GAA)_28_ and (GAA)_33_ strains without (glu, light blue) or with (gal, dark blue) galactose induction of nCas9-D10A expression. Error bars represent 95% confidence intervals. (E) Expansion rate (left) and scale (right) of WT (GAA)_33_ strains with galactose induction of nCas9-D10A expression. Left: Error bars represent 95% confidence intervals. Right: Dashed lines represent median of distribution.

To create cassettes with (GAA)_n_ repeats corresponding to the premutation and long-normal alleles, a yeast strain with the original *UR*-*(GAA)_100_*-*A3* cassette was grown on the SC Ura-plates, which favors repeat contractions, followed by PCR analysis of the repeat lengths in individual clones. Clones with 40, 33 or 28 GAA repeats were identified, and the repeat lengths were confirmed by DNA sequencing. In this approach, the unique part of the *URA3* intron was not altered, thus, the total intron lengths in the resultant cassettes were shorter compared to the original cassette. Targeted DNA nicks were introduced adjacent to the repeats in those strains and repeat expansion rates were measured as described above.

For the carrier-size (GAA)_40_ repeat, both 5’ GAA-strand and 5’ TTC-strand induction of nicks caused ∼20-fold increase in 5-FOA^r^ rate, and practically all of the 5-FOA^r^ colonies contained large-scale repeat expansions (Figure 2B&C). This was the first observation of large-scale expansions of the carrier-size repeat in any experimental system studied before by us and others^38,74^ (Figure 2C). Analysis of expansion scale demonstrated that most of the expanded repeats were several times over the size of the original (GAA)_40_ run (Figure 2C, right).

When we induced 5’-nicks at the long-normal-size (GAA)_33_ repeat, there was a smaller, but significant, 8-fold increase of in the rate of 5-FOA^r^ colonies (Figure 2D). Over two thirds of those colonies harbored expanded repeats, also exceeding the original repeat length many times over (Figure 2E). Again, large-scale expansions of long-normal repeats were never observed before.

Strikingly, there was no increase in 5-FOA^r^ rate when we induced 5’ GAA-strand nick at the normal size (GAA)_28_ repeat (Figure 2D), and no expansions were observed, even though it is only 15% shorter than (GAA)_33_ repeat.

In sum, we have demonstrated for the first time to our knowledge^38,74^, large-scale expansions of the premutation and long-normal (GAA)_n_ repeats induced by adjacent 5’ DNA nicks that resemble large-scale expansions common for human pedigrees.

### Genetic controls of nick-mediated expansions in dividing cells

Initially, we assumed that a uniform genetic mechanism might account for repeat expansions mediated by various DNA nicks. Our genetic data, however, proved us wrong. Table 1 summarizes the differences between the different types of nicks we experimented with in this project. Genetic controls of repeat expansions induced by individual DNA nicks are considered separately below.

**Table 1.** Summary of different types of nicks.

| Cassette orientation | Nick type | Cas9 type | gRNA | nick on lagging/leading strand template | nick upstream/downstream of repeat relative to replication fork |
| --- | --- | --- | --- | --- | --- |
| direct | 5' GAA | D10A | 1 | lagging | upstream |
| direct | 5' TTC | D10A | 2 | leading | downstream |
| direct | 3' GAA | H840A | 2 | lagging | downstream |
| direct | 3' TTC | H840A | 1 | leading | upstream |
| inverted | 5' GAA | D10A | 1 | leading | downstream |
| inverted | 5' TTC | D10A | 2 | lagging | upstream |

### 5’ GAA-strand nicks

Our null hypothesis was that 5’ nick-mediated repeat expansions would depend on the flap processing. Thus, we tested DNA nucleases that either cleave flaps off or affect their processing: Rad27, Sgs1, Rad1-Rad10, Mre11 and Exo1^75–80^ (reviewed in ref. 81,82). Inactivation of these proteins had no major effect on nick-mediated repeat expansions compared to the WT (Figure S2). A smaller differential increase in nick-induced expansion rate in the *rad27-2E1* mutant is likely attributed to an increased expansion rate in this mutant without any nicks^36^. Similarly, we didn’t see any role in nick-mediated repeat expansions of the Pol32, which is a subunit of Pol δ^83^ as well as Pol ζ^84^. Pol δ can displace Okazaki fragment to create flaps during lagging strand synthesis, and synthesizes long stretches of DNA in various DNA repair pathways^85,86^. Finally, inactivation of DNA helicases Chl1^87^, Srs2^88,89^, Mph1^90,91^ and Rad5^92^, or structure specific resolvases Yen1 and Mus81 had no effect on nick-mediated expansions (Figure S2).

Since we have previously demonstrated that ssDNA binding protein RPA plays an important role in GAA repeat instability^93,94^, we tested whether it also affects nick-mediated repeat expansions. We overexpressed all three subunits of yeast RPA, Rfa1, Rfa2 and Rfa3, by introducing the previously described^93^ multi-copy plasmid and confirmed overexpression by Western blotting (Figure S4). After inducing 5’ GAA-strand nick in the strain carrying the RPA-OE plasmid, we observed only a small (2-fold) decrease in the expansion rate compared to WT rate.

We then examined the role of three key genes required for homologous recombination in yeast: *RAD52*, *RAD51* and *RAD54*^95–100^ (reviewed in ref. 101). Strikingly, each of these genes’ knockouts all but eradicated the increase in the repeat expansion rate caused by the 5’ GAA-strand nicks (Figure 3A). Furthermore, Rad51 knockout dramatically reduced the increase in the repeat expansion scale upon nick induction, while Rad52 and Rad54 knockouts displayed no such increase whatsoever (Figure 3B). We therefore conclude that homologous recombination is the main player in repeat expansions mediated by 5’ GAA-strand nicks.

**Figure 3.**
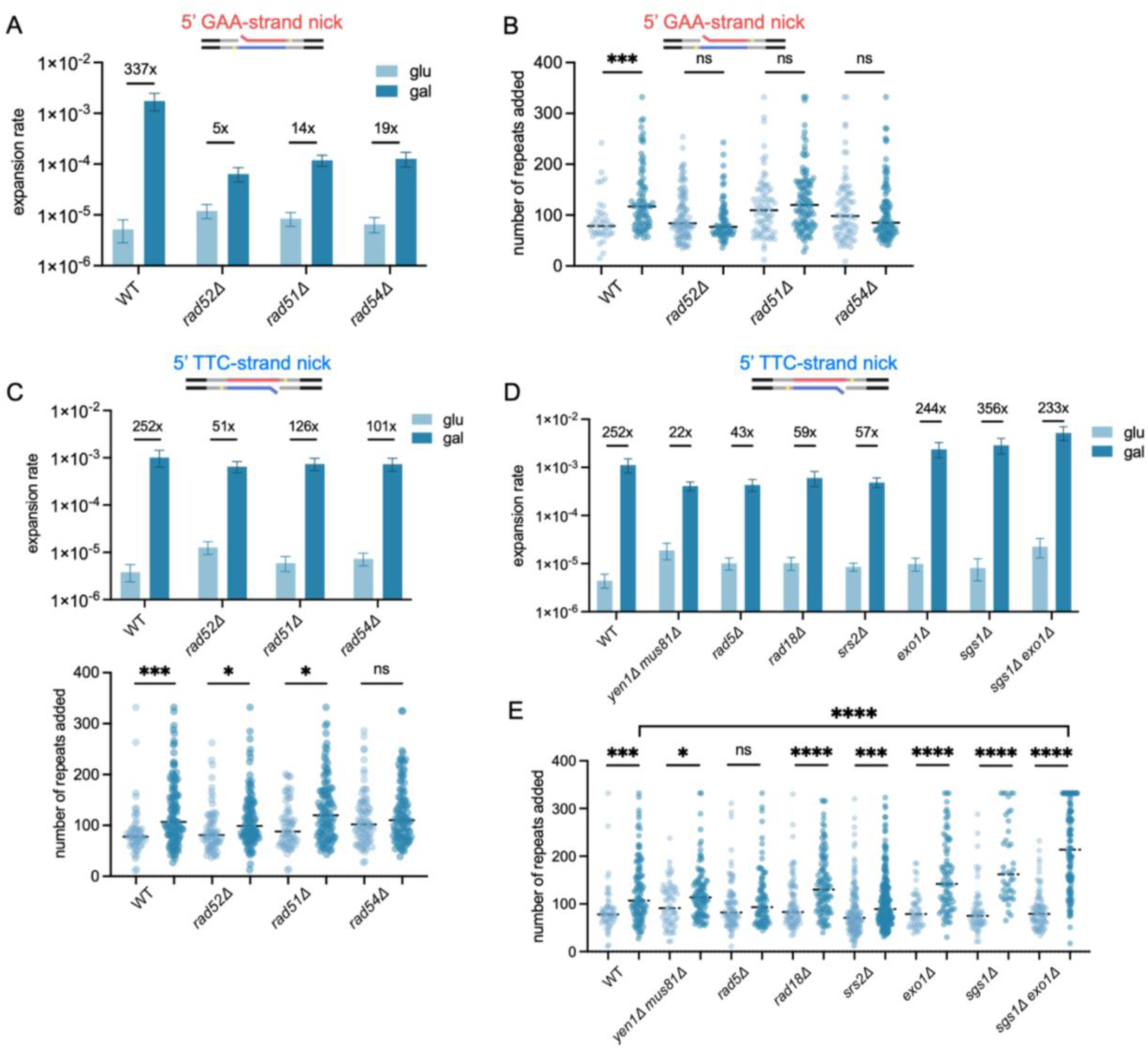
Genetic controls of the 5’ nick-mediated expansion rate and scale. (A) 5’ GAA-strand nick-mediated expansion rate of WT and *rad52Δ*, *rad51Δ* and *rad54Δ* strains without (glu, light blue) or with (gal, dark blue) galactose induction of nCas9-D10A expression. Numbers on top of each genotype show fold change of expansion rate between glucose and galactose group. Error bars represent 95% confidence intervals. (B) 5’ GAA-strand nick-mediated expansion scale of WT and *rad52Δ*, *rad51Δ* and *rad54Δ* strains without (glu, light blue) or with (gal, dark blue) galactose induction of nCas9-D10A expression. Dashed lines represent median of distribution. Statistical significance between glucose and galactose groups was calculated via Kolmogorov–Smirnov test. (C) 5’ TTC-strand nick-mediated expansion rate (top) and scale (bottom) of WT and *rad52Δ*, *rad51Δ* and *rad54Δ* strains without (glu, light blue) or with (gal, dark blue) galactose induction of nCas9-D10A expression. Top: Numbers on top of each genotype show fold change of expansion rate between glucose and galactose group. Error bars represent 95% confidence intervals. Bottom: Dashed lines represent median of distribution. Statistical significance between glucose and galactose groups was calculated via Kolmogorov–Smirnov test. (D) 5’ TTC-strand nick-mediated expansion scale of WT and various mutant strains without (glu, light blue) or with (gal, dark blue) galactose induction of nCas9-D10A expression. Expansion rate of WT and various mutant strains without (glu, light blue) or with (gal, dark blue) galactose induction of nCas9-D10A expression. Numbers on top of each genotype show fold change of expansion rate between glucose and galactose group. Error bars represent 95% confidence intervals. (E) Expansion scale of WT and various mutant strains without (glu, light blue) or with (gal, dark blue) galactose induction of nCas9-D10A expression. Dashed lines represent median of distribution. Statistical significance between glucose and galactose groups was calculated via Kolmogorov–Smirnov test.

### 5’ TTC-strand nicks

Since the fold increase in the rate of 5’ TTC-strand nick-mediated expansions was similar to that of 5’ GAA-strand nick, we initially hypothesized that they share a similar HR-based mechanism. Contrary to these expectations, deletions of *RAD52*, *RAD51* or *RAD54* genes did not significantly alter expansion rate as compared to the WT (Figure 3C). Deletions of *POL32* and *REV3* genes, or RPA overexpression did not affect repeat expansions triggered by 5’ TTC-strand nicks (Figure S3). Deletions of individual *YEN1* or *MUS81* genes, which encode structure-specific endonucleases involved in resolution of Holliday junctions^102,103^ did not have an effect, while simultaneous deletion of both genes significantly rescued nick-mediated expansions (Figure 3D). The deletion of *RAD5*, a gene encoding a multifunctional protein involved in post-replication repair (PRR) and template switching^92,104^ resulted in a similar decrease in nick-mediated expansions. The deletion of *RAD18*, the gene encoding a protein involved in PRR^105^ led to smaller decrease in nick-mediated expansion rate, which was border-line significant (Figure 3D). Deletion of the *SRS2* gene, encoding DNA helicase and DNA-dependent ATPase involved in DNA repair and recombination^88,89,106^, resulted in a small but reproducible decrease in nick-mediated expansions (Figure 3D). Deletion of the *MPH1* gene, encoding another DNA helicase^90,91^, did not significantly affect expansion rate (Figure S3).

When either *EXO1* or *SGS1,* genes involved in the long-range resection of DSBs and SSBs^76,80^, were deleted individually, we observed small increases in 5’ TTC-strand nick-mediated expansions that were borderline significant (Figure 3D). The deletion of both genes together led to a significant, 4-fold increase in the nick-mediated expansion over the WT (Figure 3D). Furthermore, the scale of the expansions also dramatically increased (Figure 3E), with over 20% of expanded repeats being out of the range of the ladder we used for our gel electrophoresis which was 1350 bp (∼332 repeats added).

### 5’ nicks with the inverted cassette

We reasoned that the difference in genetic mechanism between 5’ GAA-strand nick and 5’ TTC-strand nick could be due to the position during DNA replication, with the former interrupting the lagging strand template and the latter interrupting the leading strand template. To explore this possibility, we inverted our cassette relative to the ARS306. Since the entire cassette was inverted, the repeat tract was in the opposite orientation relative to replication but not transcription (Figure 4A). A small repeat contraction (95 instead of 100 repeats) occurred in the process of strain construction, but it did not significantly alter the expansion rate.

**Figure 4.**
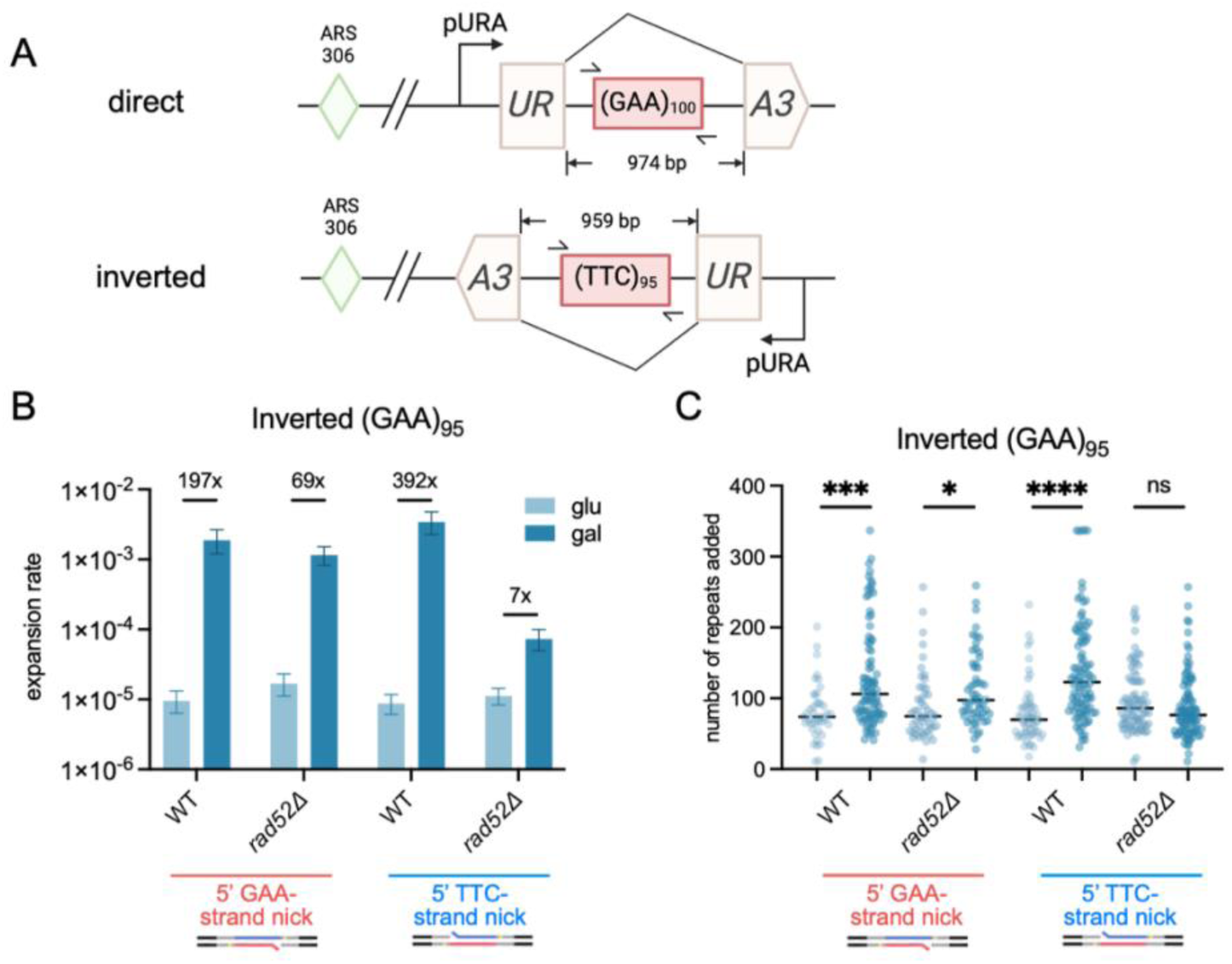
Expansion of WT and *rad52Δ* strains with the inverted cassette. (A) Schematic of the direct (top) and inverted (bottom) cassettes. In the inverted cassette, the entire *URA3* gene is flipped relative to the origin of replication ARS306, resulting in the GAA strand being the leading strand template. The 5’ GAA-strand nick is downstream, and the 5’ TTC-strand nick is upstream of the repeat relative to the replication fork direction. (B) Expansion rate of WT and *rad52Δ* strains without (glu, light blue) or with (gal, dark blue) galactose induction of nCas9-D10A expression. Numbers on top of each genotype show fold change of expansion rate between glucose and galactose group. Error bars represent 95% confidence intervals. (C) Expansion scale of WT and *rad52Δ* strains without (glu, light blue) or with (gal, dark blue) galactose induction of nCas9-D10A expression. Dashed lines represent median of distribution. Statistical significance between glucose and galactose groups was calculated via Kolmogorov– Smirnov test.

Upon cassette inversion, the same 5’ GAA-strand and TTC-strand nicks interrupt the leading and lagging strand templates, respectively. Both of them increased repeat expansion rates and scales similarly to what was observed for the original cassette (Figure 4B&C). At the same time, the effect of HR on nick-mediated expansions was reversed; Rad52 knockout did not affect expansions induced by the 5’ GAA-strand nick but rescued the effect of the 5’ TTC-strand nick (Figure 4B). We conclude, therefore, that the difference in genetic controls of expansions between the 5’ GAA-strand and TTC-strand nicks was due to their positioning in the replicon.

### 3’ nicks

Given that deletions of *RAD52* or *RAD51* genes rescued expansions induced by the 5’ GAA-strand nick, we tested if the same is true for 3’ nicks. Repeat expansions induced by either of 3’ nicks were indeed fully rescued by either gene knockouts (Figure 5A&B). Also, the scale of the expansions decreased to the WT level in both mutants (Figure 5A&B).

**Figure 5.**
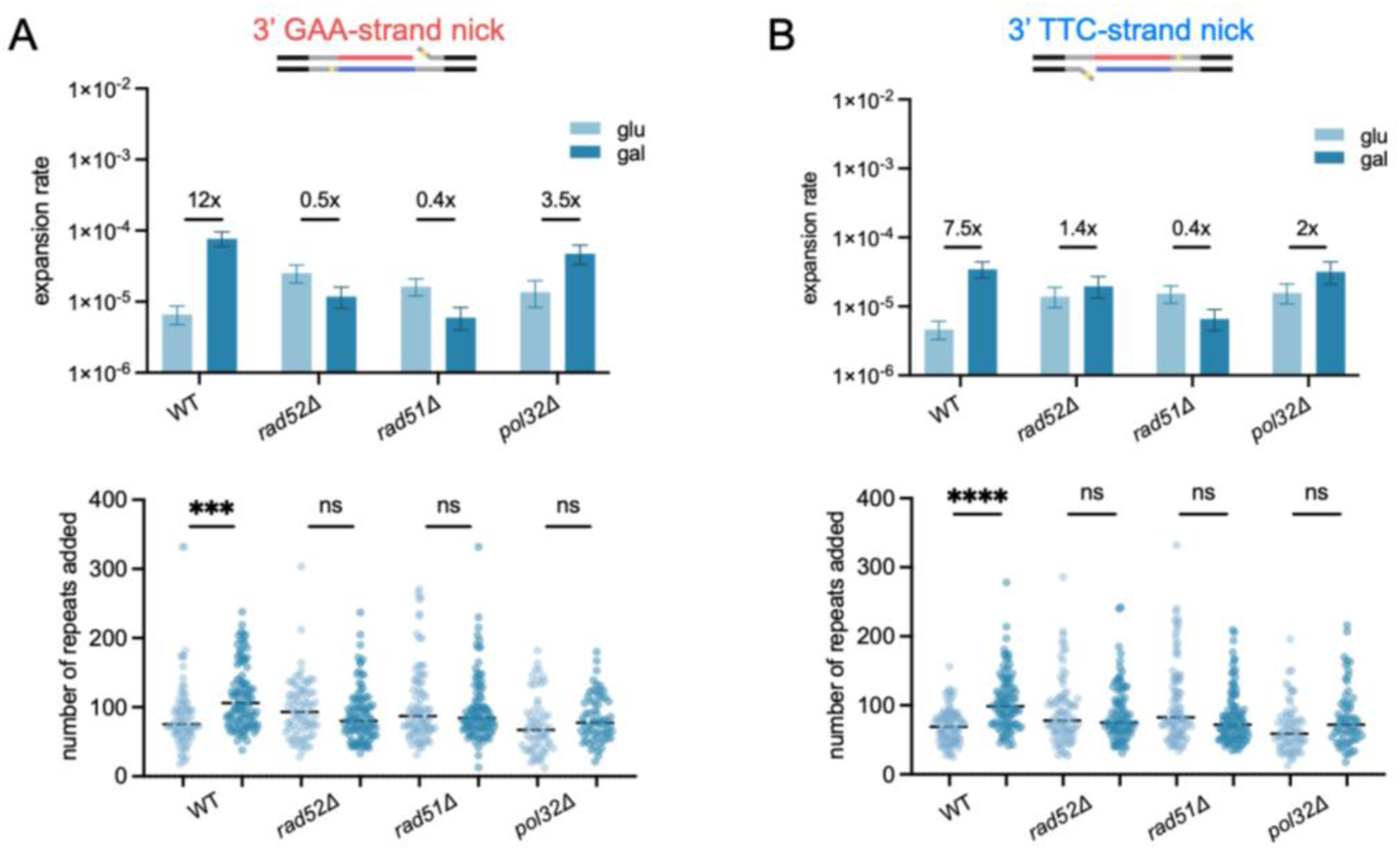
Genetic controls of the 3’ nick-mediated expansion rate and scale. (A) 3’ GAA-strand nick-mediated expansion rate (top) and scale (bottom) of WT, *rad52Δ*, *rad51Δ* and *pol32Δ* strains without (glu, light blue) or with (gal, dark blue) galactose induction of nCas9-H840A expression. Top: Numbers on top of each genotype show fold change of expansion rate between glucose and galactose group. Error bars represent 95% confidence intervals. Bottom: Dashed lines represent median of distribution. Statistical significance between glucose and galactose groups was calculated via Kolmogorov–Smirnov test. (B) 3’ TTC-strand nick-mediated expansion rate (top) and scale (bottom) of WT and *rad52Δ*, *rad51Δ* and *pol32Δ* strains without (glu, light blue) or with (gal, dark blue) galactose induction of nCas9-H840A expression. Top: Numbers on top of each genotype show fold change of expansion rate between glucose and galactose group. Error bars represent 95% confidence intervals. Bottom: Dashed lines represent median of distribution. Statistical significance between glucose and galactose groups was calculated via Kolmogorov–Smirnov test.

We then examined the role of Pol32, the processive subunit of Pol δ, which is a key player in break-induced replication (BIR) but not in other recombination pathways^107^. Deletion of the *POL32* gene partially rescued the rate of 3’ nick-mediated expansions (Figure 5A&B). Furthermore, the scale of expansions in the *pol32Δ* strain was much smaller. We conclude that Pol32 protein is involved in 3’ nick-mediated expansion. Since Rad52, Rad51 and Pol32 proteins are all involved in BIR, we believe that 3’ nick-mediated repeat expansions likely occur via BIR. This suggestion is in agreement with a recent study showing the role of BIR in DNA nick repair in fission yeast^108^.

### Cas9 dead protein bound downstream of the GAA repeat induces its expansions

To control for the effect of the Cas9 protein binding per se on GAA repeat expansions, we conducted the same experiments using a plasmid expressing the CAS9 gene with mutations in both nuclease domains, termed Cas9 dead (dCas9). dCas9 was bound either upstream (when guided with gRNA1) or downstream (when guided with gRNA2) of the repeat relative to the replication direction (Figure 6A). The presence of dCas9 upstream of the repeat did not significantly affect its expansion rate or scale (Figure 6B). When positioned downstream of the repeat, dCas9 increased its expansion rate 15-fold (Figure 1E) and significantly increased expansion scale (Figure 6B).

**Figure 6.**
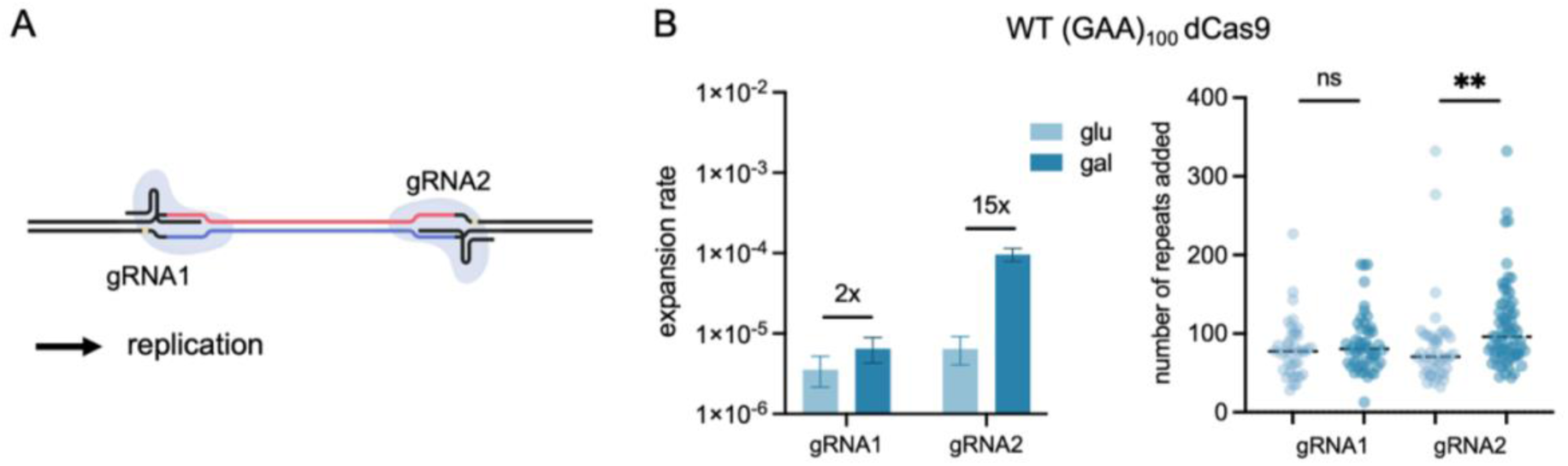
Effect of dCas9 on (GAA)_100_ repeat expansion. (A) Schematic showing positioning of dCas9 bound to gRNA1 or gRNA2 on DNA. Replication from ARS306 goes from left to right. (B) Expansion rate (left) and scale (right) of WT strains carrying combinations of dCas9 plasmid and gRNA1 or gRNA2 plasmid without (glu, light blue) or with (gal, dark blue) galactose induction of Cas9 expression. Left: Numbers on top of each genotype show fold change of expansion rate between glucose and galactose group. Error bars represent 95% confidence intervals. Right: Dashed lines represent median of distribution. Statistical significance between glucose and galactose groups was calculated via Kolmogorov–Smirnov test.

We speculate that these increases could be due to stalling of the replication fork upon encountering tightly bound dCas9 protein. When replication machinery encounters dCas9 upstream of the repeat, its stability should not be affected by the fork stalling. In contrast, when replisome stalls downstream of the repeat, fork stalling and restart can drive repeat instability. Supporting this hypothesis, knockout of the Rad5 protein, involved in PRR and fork restart, leads to a decrease in the dCas9 mediated repeat expansions (Figure S5).

### 5’ GAA-strand nicks induce expansions in non-dividing yeast cells

Repeat expansions commonly occur in post-mitotic somatic cells^109^. Our previous study showed that (GAA)_100_ repeat undergo expansions in quiescent yeast cells (Q cells)^39^, *i.e.* in cell cells arrested in G0 phase. Thus, we were interested in whether our nickases could induce expansions in Q cells as well. Quiescence was induced similarly as previously described^39^, with minimal media modification to assure maintenance of plasmids expressing the nickase system (Table S4). Yeast was cultured in phosphate-limited -LEU +Hyg media and then transferred into -LEU +Hyg media with less sugar and no phosphate. For phosphate-limited media, only glucose was used; for no-phosphate media, either glucose or galactose was used. Fluctuation tests were performed with cells 15 minutes after transition from phosphate limited media to no phosphate media (QD0) as well as 6 days after transition (QD6) (Figure 7A). Since quiescent cells do not divide, this approach allows us to measure the frequencies of repeat expansions rather than their rates.

**Figure 7.**
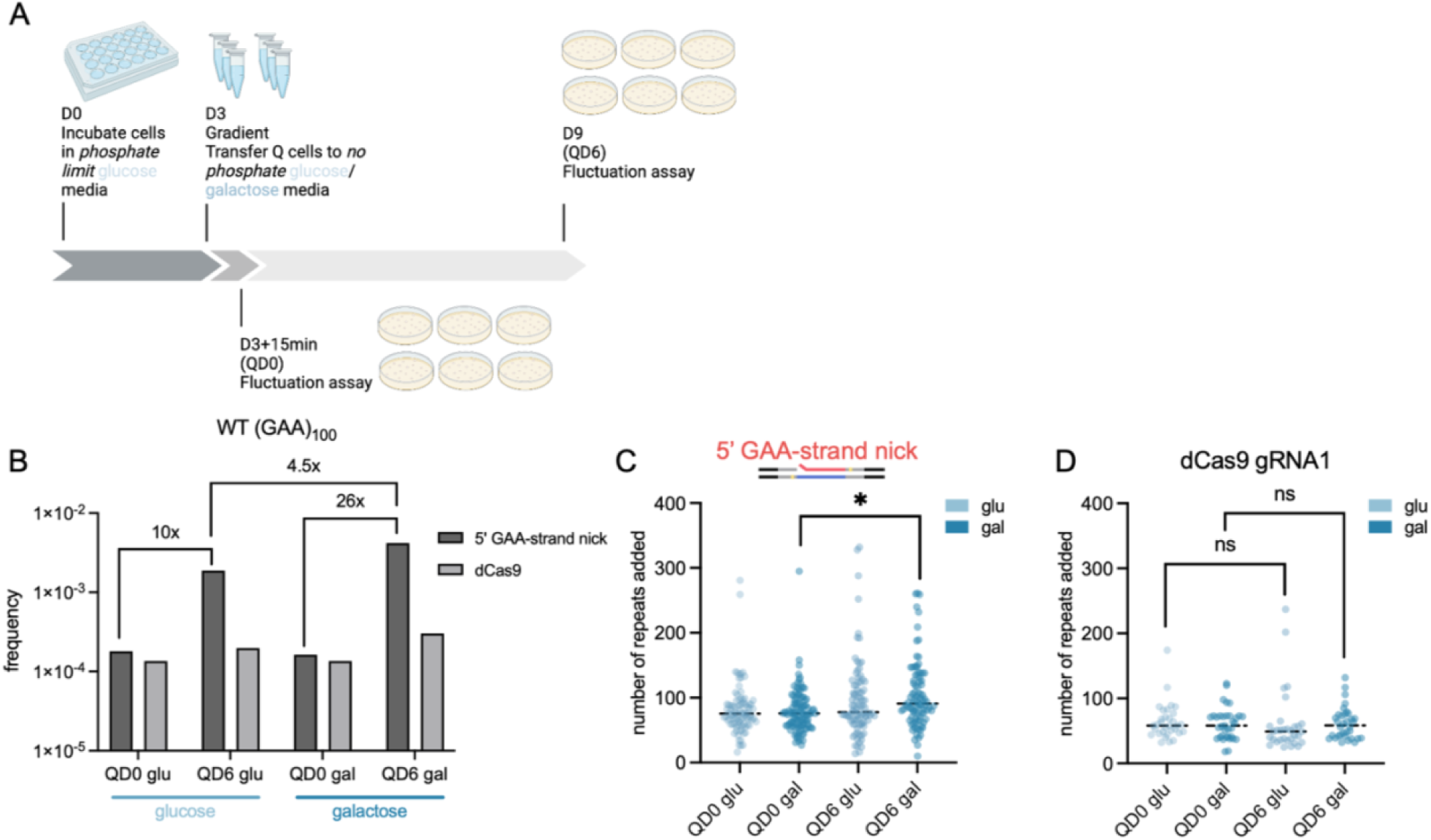
5’ GAA-strand nick-mediated expansion in quiescent yeast cells. (A) Schematic of experimental setup for inducing quiescence and fluctuation tests. Quiescence was induced by limiting phosphate concentration in liquid media. (B) Expansion frequency of quiescent WT without (glu, light blue) or with (gal, dark blue) galactose induction of Cas9 expression. Dark grey: strains carrying nCas9-D10A and gRNA1 plasmids. Light grey: strains carrying dCas9 and gRNA1 plasmids. (C) Expansion scale of quiescent WT strains without (glu, light blue) or with (gal, dark blue) galactose induction of nCas9-D10A expression. Dashed lines represent median of distribution. Statistical significance between QD0 and QD6 was calculated via Kolmogorov–Smirnov test. (D) Expansion scale of quiescent WT strains carrying of dCas9 and gRNA1 plasmids without (glu, light blue) or with (gal, dark blue) galactose induction of Cas9 expression. Dashed lines represent median of distribution. Statistical significance between QD0 and QD6 groups was calculated via Kolmogorov–Smirnov test.

After 6 days in quiescence, expansion frequency of cells in galactose media increased by 26-fold compared to QD0 (Figure 7B). Unexpectedly, however, expansion frequency increased 10-fold even in glucose-containing media (Figure 7B). At the same time, expansion scale was significantly changed during chronological aging of cells cultivated in galactose-containing media (Figure 7C). Altogether, our data indicate that 5’ GAA-strand nicks lead to increased repeat expansions in non-dividing cells as well.

One explanation for increased expansion frequency in glucose-containing media could come from the leakiness of the *GAL1* promoter in the presence of glucose. In dividing cells, *GAL1* promoter is strongly repressed in the presence of glucose owing to the inactivity of the Gal4 activator combined with the binding of Mig repressors. Given that Q cells are very different from dividing cells^110^, it is foreseeable that the *GAL1* promoter may not be fully repressed by glucose, resulting in residual expression of the nCas9 protein.

To address this possibility, we induced quiescence in cells carrying dCas9 and gRNA1 plasmids. In this case, we observed only a minimal, 2-fold increase in the repeat expansion frequency both in glucose- and galactose-containing media (Figure 7B) and no significant increase in the expansion scale (Figure 7D). The frequencies of repeat expansions in galactose-containing media differ ∼30-fold between nCas9 and dCas9, indicative of a strong elevation of repeat expansions in quiescent cells upon 5’-nick induction.

Taken together, we show that nicks induce massive expansions of GAA repeats in both dividing and non-dividing cells. In dividing cells, the position of the GAA tract relative to the replication fork plays major role in determining the mechanism of such expansion.

## Discussion

In this study, we demonstrated that DNA nicks introduce adjacent to the (GAA)_n_ repeat promote its large-scale expansions in our intronic yeast system. Most strikingly, DNA nicks 5’ to the repeat led to an over-two-orders-of-magnitude increase in the rate of very large-scale expansions for the pathogenic-size (GAA)_100_ repeat (Figure 1C&D).

The large scale of this effect led us to hypothesize that 5’ nicks might also trigger massive expansions of the premutation (GAA)_40_, as well as the long-normal (GAA)_33_ repeats, which indeed turned out to be the case (Figure 2). To the best of our knowledge, this is the first demonstration of large-scale expansions of the premutation and long-normal repeats in any experimental system studied. This is an important observation because, while premutation alleles do expand during intergenerational transmission in humans, the mechanisms of these events are not well understood. We understand even less about how long-normal size alleles convert into the premutation alleles. Early models of repeat expansions suggested that they occur via multiple rounds of small strand slippage events during replication and/or repair^111^ events. Based on our data, it is tempting to propose an alternative mechanism for repeat expansions, in which DNA nicks originated at a repeat in rapidly replicating human pre-meiotic cells serve as a trigger of large-scale expansions. Similar to the ‘toxic oxidation cycle’^53^, more nicks could be generated as the repeat grows larger, leading to massive expansions observed in human pedigrees. In our case, this is particularly evident for the premutation- and long-normal-size repeats, where expansions are several folds longer than original repeats (Figure 2C&E).

Another intriguing observation is that the threshold of nick-mediated expansions in our intronic yeast system stands somewhere between 28 and 33 repeats, *i.e.* right at the border between normal and premutation alleles in human pedigrees. This could be attributed to the stability of a triplex DNA structure formed by the GAA repeat, which increases exponentially with the repeat’s length^112,113^.

Finally, we found that 5’ GAA-strand nicks also increase expansion frequency in non-dividing yeast cells, albeit to a lesser extent that in dividing cells (Figure 7B). This is an equally important observation, since GAA repeat expansions in post-mitotic cells appear to contribute to the FRDA progression^23,29^. A recent study showed that there is a high-level accumulation of DNA nicks in post-mitotic neurons^114^. Considering that the nervous system is the most severely affected in REDs, it is plausible to speculate that repair of DNA nicks could play an important role in repeat expansions in neurons as well as other metabolically active post-mitotic cells.

Our genetic analysis revealed that most types of nick-mediated expansions in our system involved recombination proteins Rad51 and Rad52. Expansions driven by the 5’ GAA-strand were highly dependent on Rad52, Rad51 and Rad54 proteins (Figure 3A&B), implicating canonical homologous recombinational pathway of DSB repair. Expansions caused by either 3’ nick were also strongly dependent on Rad51 and Rad52, and required Pol32 (Figure 5), pointing to another pathway of DSB repair: break-induced replication (BIR).

To our surprise, however, genetic controls of expansions mediated by the 5’ TTC-strand nick appeared to be quite different: *RAD52* and *RAD51* genes had no bearing, deletions of *RAD5, MUS81, YEN1,* and *SRS2* genes each led to a partial rescue of repeat expansions, while deletions of *SGS1* and *EXO1* genes increased expansion rate (Figure 3D). We hypothesized that the difference in genetic controls between the 5’ GAA-strand and 5’ TTC-strand nicks depends on which strand is being interrupted during DNA replication: the lagging strand template by the former and the leading strand template by the latter. This hypothesis was proven correct, since inverting our *UR*-*(GAA)_100_*-*A3* cassette relative to the replication origin flipped genetic requirements for expansions generated by the two nicks. It is also in an agreement with the earlier data showing that nicks in the lagging, but not in the leading strand template, are preferentially repaired by HR^115^.

A DNA nick in the lagging strand template can either lead to a fork collapse^116^ or be bypassed by the replisome, leaving behind a two-ended DSB^117^, which can be repaired by sister-chromatid exchange^118^. A DNA nick in the leading strand template, in contrast, cannot be bypassed because the CMG helicase moving along the leading strand template would simply slide off ^116,119–121^, resulting in fork collapse and the formation of a one-ended DSB with no sister chromatid to rescue. This consideration led us to propose working models for the 5’ nick-mediated (GAA)_n_ repeat expansion in dividing cells. During replication, the 5’ GAA-strand nick is converted into a two-ended DSB. This DSB could be repaired by the synthesis-dependent strand-annealing (SDSA) pathway of HR using the sister chromatid as repair template. In course of this process, the newly synthesized repetitive strand could misalign with the donor chromatid resulting in expansion (Figure 8 left). The 5’ TTC-strand nick, in turn, leads to fork collapse and the formation of a seemingly unrepairable one-ended DSB. This situation could, however, be rescued by the converging replication fork for which this nick will break the lagging strand template upstream of the repeat. Technically, this would lead to over-replication of the repeat and a version of DSB repair via Rad5-mediated template-switching^122^ leading to repeat expansions (Figure 8 right).

**Figure 8.**
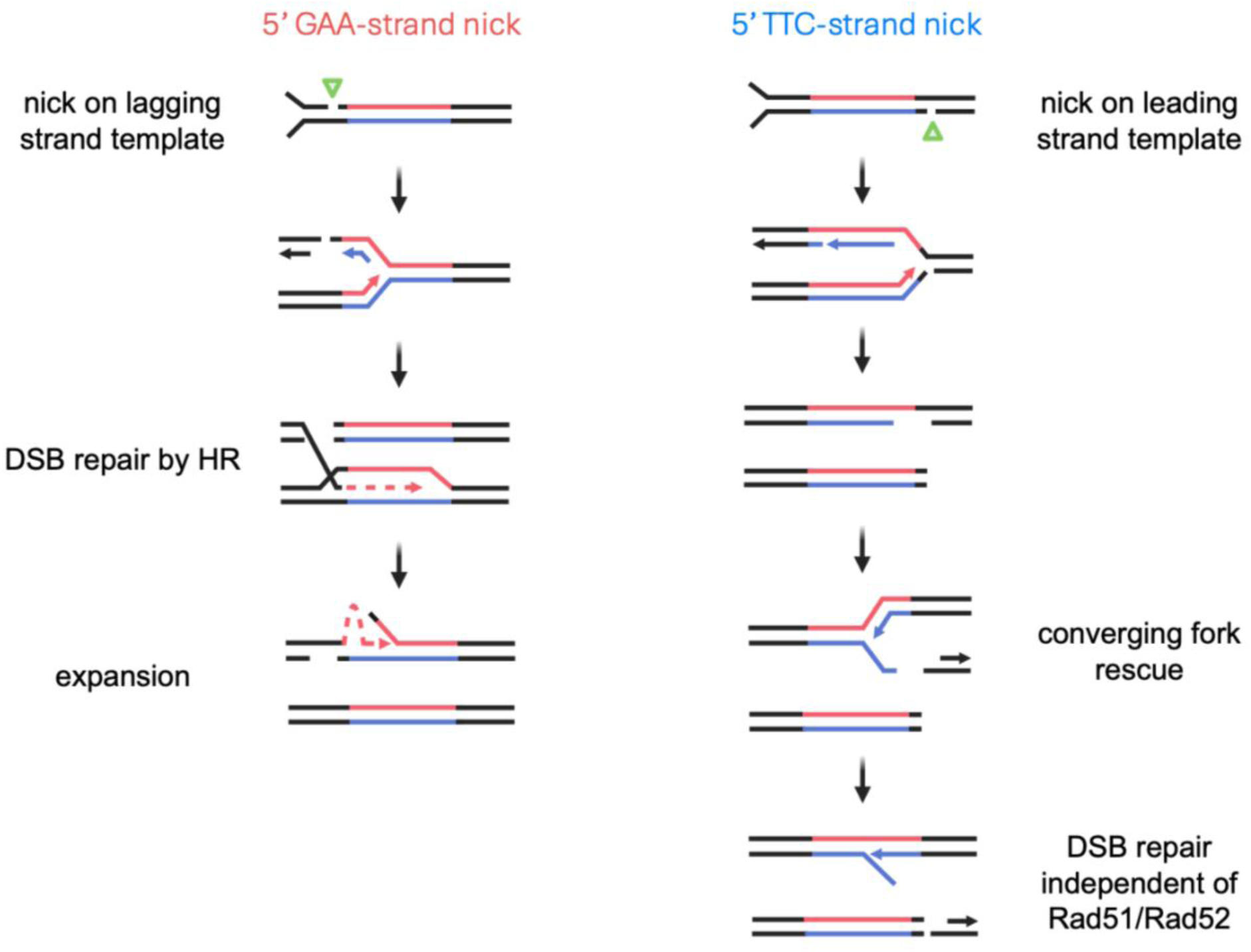
Model of 5’ nick-mediated (GAA)_n_ repeat expansion. Left: The 5’ GAA-strand nick on the lagging strand template is converted into a two-ended DSB during replication, which is repaired by HR. Expansion occurs through misalignment of the repeat tract during strand invasion. Right: The 5’ TTC-strand nick on the leading strand template is converted into a one-ended DSB during replication when the fork collapses due to CMG helicase falling off. Sister chromatids are held together by cohesin. The one-ended DSB is rescued by a converging fork and repaired by an unknow mechanism independent of Rad51/Rad52. Expansion could occur via misalignment or flap displacement.

Alternatively, the nicks could be efficiently induced immediately after replication at nucleosome free DNA segments. Nicks in nascent strands could be biased towards repair by ligation^51,123^. However, nicks in the leading or lagging strand templates might not get as effectively ligated. 5’ GAA-strand nick in the lagging strand template could be converted into a DSB before the Okazaki fragment is maturated and repaired by HR similarly to a model described above (Figure S6 left). 5’ TTC-strand nick in the leading strand template could be repaired by template switching using the complementary single-stranded lagging strand template, – a process, which would not require Rad51 (Figure S6 right).

While these are plausible hypotheses, we are still uncertain why none of the genes we studied has a major effect on repeat expansions driven by the 5’ TTC-strand nick mediated. One possibility could be that those expansions accumulate over the course of several genetic processes. This idea is supported by the fact that even catalytically dead dCas9 bound at the 3’-end of the repeat cause a significant increase in repeat expansions. Thus, expansion mediated by the 5’ TTC-strand nick could be at least partially attributed to protein-mediated fork stalling.

A recent study showed that nCas9-H840A is capable of inducing DSBs with low efficiency^124^. Since we used nCas9-D10A for both 5’ nicks and H840A for both 3’ nicks (Figure S1C), this observation does not affect our conclusions for the 5’ nicks. However, this might explain the difference in both nick-mediated expansion rate and mechanisms between 5’ and 3’ nicks: While Rad52 and Rad51 proteins are both involved in highly elevated expansions caused by the 5’ GAA-strand nick and a more modest increase by 3’ nicks, Pol32 protein was only implicated in expansions caused by 3’ nicks. Therefore, one possibility, is that the moderate increase in repeat expansions caused by 3’ nicks could be due to DSBs introduced with low efficiency by nCas9-H840A. Alternatively, the difference in genetic controls between the 5’ GAA-strand and 3’ nicks could be grounded in differences in DNA binding between nCas9-H840A and nCas9-D10A. Their DNA binding is not symmetrical, as there is a target strand which is complementary to gRNA and a non-target strand^125^. Since nCas9-H840A nicks the non-target strand, its 3’-end is free to move out of the nuclease domain making it accessible for nick repair pathway, which means that it is much less likely be converted into a DSB during replication. If we assume that nCas9-H840A indeed induces nicks at 3’ of the repeat tract, the genetic controls of 3’ nick-induced expansions support a model that the expansions were mediated by BIR (Figure S7).

We showed that 5’ GAA-strand nicks also increased expansion frequency in quiescent cells. Obviously, nick-mediated repeat expansions in Q-cells cannot rely on DNA replication. We have previously shown potent recombinational events triggered by GAA repeats during the G1-phase of the cell cycle^44^ and in Q cells^39^. Others have demonstrated that recombination can occur during G1-phase in mammalian cells^126^. It is therefore possible that repeat expansions triggered by the 5’ GAA-strand nick involve a recombinational pathway yet to be characterized.

Very recently, it was found that targeted DNA nicks as well as DSBs led to increased instability of the G_4_C_2_ repeat, which is associated with ALS and FTD in a transgenic mouse model^68^. The authors proposed that nick-induced repeat expansions involve HR^68^. Note, however, that they did not observe any increase in expansions or contractions of GAA repeats^68^. In contrast, we observed dramatic increases in GAA repeat expansions driven by targeted nicks in our yeast system. Further studies are needed to understand the reasons for this dramatic difference in the outcome of DNA nicks on GAA repeat expansions in mice and yeast.

## Methods

### Plasmid construction

To generate the inverted cassette, the *pURA-UR-(GAA)_100_-A3-TRP1* cassette was amplified with primers FII_A3_CCCTT84_UR_LR and FII_A3_CCCTT84_UR_LF, and the vector backbone was amplified with primers FI_pJH21backbone_LF and FI_pJH21backbone_LR. These primers contain homology of each other. Fragments were then ligated with Gibson assembly (NEBuilder® HiFi DNA Assembly Master Mix #E2621L, NEB).

To create the pJA29_pGal_Cas9dead plasmid, we amplified a fragment of the Cas9 gene containing the H840A mutation from the pRS415_pGal-nCas9(H840A) plasmid (Addgene #79616) with primers NcoI_Cas9_F and Cas9_R. We then digested this fragment with the *Sbf*I and *Asc*I restriction enzymes and ligated with a fragment from the pRS415_pGal-nCas9(D10A) plasmid (Addgene #79618) digested with the same pair of enzymes. The resulting plasmid is identical to the pRS415 plasmids except it contains a catalytically dead Cas9 with both D10A and H840A mutations.

To create the gRNA plasmid backbone called pSK26_gRNA_Hyg we digested the bRA89 plasmid (Addgene #100950) with the *Ngo*MIV and *Cla*I restriction enzymes and ligated the 5736 bp fragment onto itself by inserting a short filler fragment digested with the same pair of enzymes. To insert specific gRNA sequences into the pSK26_gRNA_Hyg, primers containing the gRNA sequence and the *Bpl*I restriction enzyme recognition sites were ligated with the backbone cut by *Bpl*I.

Sequences of all the plasmids used in this study were sequenced by Sanger sequencing (Genewiz) or long read sequencing (Plasmidsaurus and Genewiz) and their full sequences are available in the supplemental materials.

### Yeast strain construction

All yeast strains used in this study were constructed of the CH1585 derivative SMY923 (*MAT**a**, leu2Δ1, trp1Δ63, ura3Δ, his3Δ200, MIP1, HAP1*). The integration of the *pURA-UR-(GAA)_100_-A3-TRP1* was previously described in ref. 37. To obtain strains with premutation or long-normal GAA alleles, cells with (GAA)_100_ allele were diluted and plated on SC -Ura media to select for natural contractions. Colony PCR was performed to analyze repeat length and colonies with desired repeat length were used for the experiments. Mutants were made using either direct gene replacement or a simplified CRISPR-Cas9 approach^127^. Knockout transformants were confirmed by PCR using both internal and external primers against the targeted gene (Table S2). Point mutations were confirmed by Sanger sequencing (Genewiz).

To express the nickase system, the combination of a plasmid with a Cas9 nickase (pRS415_pGal-nCas9(D10A) or pRS415_pGal-nCas9(H840A)) or a dead Cas9 (pJA29-pGal-Cas9dead), and a plasmid with a guide RNA under the SNR52 promoter (pLZ1_GAA_gRNA1_HYG or pLZ2_GAA_gRNA2_HYG) were co-transformed at 300 ng each into yeast. The Cas9 plasmids use the LEU2 gene as and the gRNA plasmids use the HYG gene as selection markers, thus we maintained the transformants on proline rich media with Hygromycin B and without leucine to facilitate plasmid retention.

### Fluctuation tests

Fluctuation test protocol was adapted from ref. 69. In short, cells were diluted and plated on proline rich media without leucine containing Hygromycin B, with either glucose or galactose for 3 to 5 days at 30 °C until colonies of desired size are formed. Individual colonies were dissolved in 200 μL sterile water, serially diluted, and plated on YPD or 5-FOA media (0.095% 5-FOA). Genomic DNA of each colony plated was extracted and PCR was performed to confirm the repeat tract length was not altered. Colonies with altered repeat tract were excluded from analysis. For each genotype, 8 to 16 colonies were used to perform fluctuation tests. Colonies grown on selective (5-FOA) or non-selective (YPD) were counted after 3 days of incubation at 30 °C. Genomic DNA was extracted from colonies grown on 5-FOA media and repeat lengths were analyzed via PCR. Expansion rates or frequencies were calculated based on number of expanded colonies and total cell count from non-selective media using *FluCalc* software^69^.

### Western blotting

Strains were grown in -His dropout media to log phase (OD_600_=0.8), protein extraction (5OD) was conducted as described in ref. 128. Total protein extract was separated on SDS-PAGE gel, stained with Ponceau S staining (Fisher BioReagents #BP103-10) for total protein normalization, and probed with an anti-RFA antibody (Agrisera AS07 214). BenchMark™ Pre-stained Protein Ladder (Invitrogen #10748010) was used.

### GAA repeat expansion distribution

To build the distribution of GAA repeats added, colonies were randomly selected from each of the 5-FOA plate. Numbers of colonies selected from each plate were calculated roughly based on the ratio of total colonies grown on said plate to total colonies grown on all plates from the same assay for each genotype. Genomic DNA was extracted with lyticase (Sigma-Aldrich L4025), and PCR was performed with A2 and B2 or UC1 and UC6 primers to amplify GAA repeat tract. PCR products were run on 1.5% agarose gel alongside 50-bp DNA ladders (NEB #N0556S). *Image Lab* software (Bio-Rad) was used to estimate the size of the repeat tract. If two or more colonies from the same 5-FOA plate showed identical repeat size, that size is only counted once in the distribution.

### Quiescent cell experiment

Quiescent cell mutation frequency assay was adapted from ref. ^39^. Phosphate-limited media or no phosphate media was made with yeast nitrogen base (YNB) without ammonium sulfate, drop-out mix minus leucine, and Hygromycin B was added to facilitate plasmid retention (media recipes in Table S4). To control the expression of nCas9 or dCas9 protein, galactose or glucose was used in no phosphate media. Around 10 cells were seeded in 1.2 mL phosphate-limited cultures.

### Statistical analysis

For expansion rates, 95% confidence intervals were calculated using the *FluCalc* software^69^. Difference between expansion rates with non-overlapping 95% confidence intervals were considered significant. Statistical significance of repeat length distribution was calculated by Kolmogorov–Smirnov test. The significance was defined as follows: ns, p > 0.05; *, p < 0.05; **, p < 0.01; ***, p < 0.001; ****, p < 0.0001.

## Supporting information

Supplemental Materials for Li et al, 2024

## Acknowledgements

We thank Catherine Freudenreich, Mitch McVey for helpful discussions; Elina Radchenko for constructing the SMY923 strain; Julia Hisey for helping with quiescent cell experiments; Conor Moore for helping with RPA-OE strain construction; Chiara Masnovo for helping with Western blotting.

## Author contributions

L.L., A.N.K. and S.M.M. designed research; L.L. and W.S.S. performed research; A.N.K and J.F.A. constructed Cas9 and gRNA plasmids; L.L., W.S.S. and S.M.M analyzed data; and L.L. and S.M.M. wrote the manuscript.

