## Supplemental Materials for Li et al, 2024 for "DNA Nicks Drive Massive Expansions of (GAA)_n_ Repeats"

**This PDF file includes:**

Figures S1 to S7

Tables S1 to S4

#### Supplementary figures

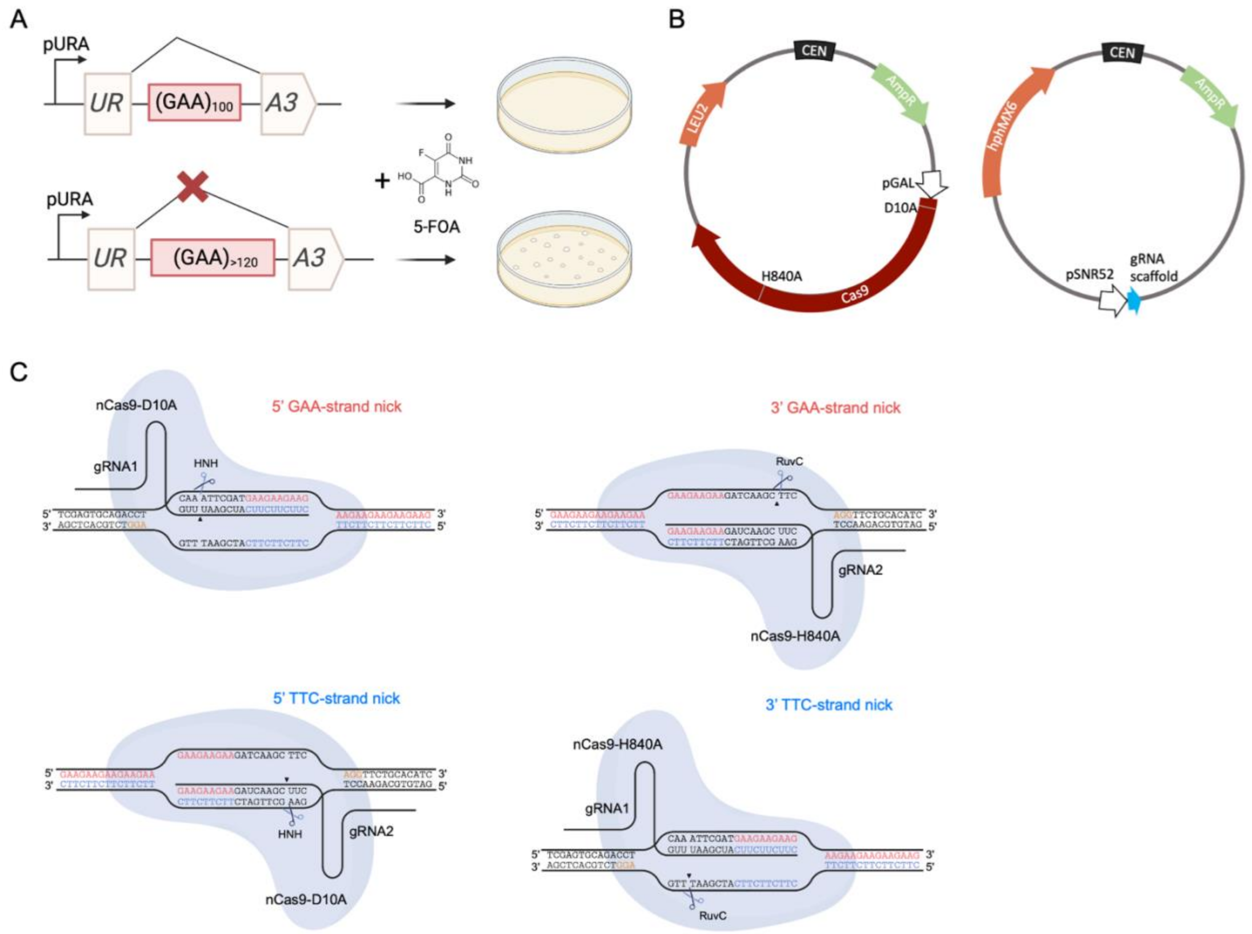

**Figure S1. Schematics of fluctuation test and the nickase system.** (A) With the $(GAA)_{100}$ repeat tract, the artificial intron of the *URA3* is spliced out and the cells are Ura+ and sensitive to 5-FOA. When the $(GAA)_n$ repeat undergoes expansion of over ~20 repeat units, the intron cannot be spliced out during transcription, resulting in inactivation of the *URA3* gene. Cells then become resistant to 5-FOA. (B) Schematic of nickase plasmids. Left, plasmid expressing Cas9 under the *GAL1* promoter, with either D10A, H840A or both mutations, with *LEU2* as selectable marker. Right, plasmid expressing gRNA under the *SNR52* promoter, with *hphMX6* as selectable marker.

(C) All four nick sites with flanking DNA sequence. Both 5' GAA and TTC-strand nicks are induced by nCas9-D10A while both 3' GAA and TTC-strand nicks are induced by nCas9-H840A. The homopurine GAA repeat sequence is shown in red and the homopyrimidine TTC repeat sequence. Non-repeat sequence is shown in black.

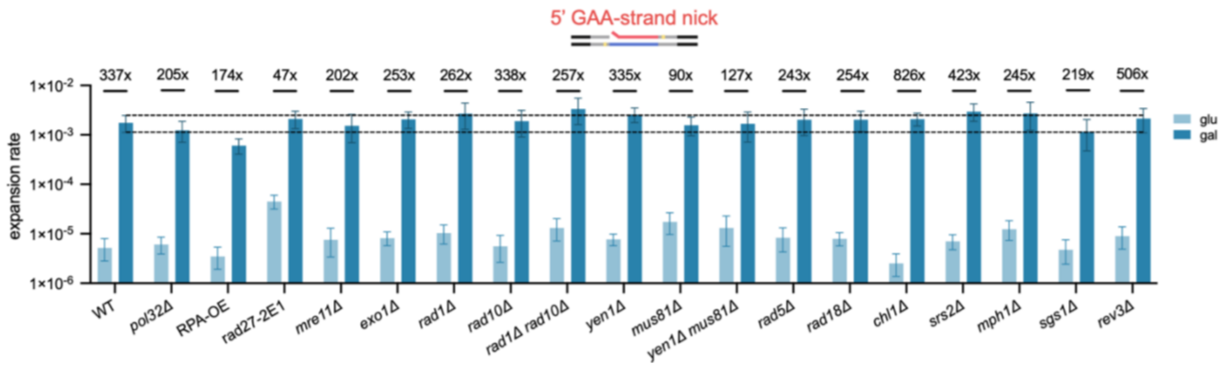

**Figure S2. 5' GAA-strand nick-induced expansion rate of additional genetic controls.** Effect of various mutants on expansion rate of without (glu, light blue) or with (gal, dark blue) galactose induced nCas9-D10A expression. Numbers on top of each genotype show fold change of expansion rate between glucose and galactose group. Error bars represent 95% confidence intervals. Dashed lines represent 95% confidence interval of WT with 5' G strand nick.

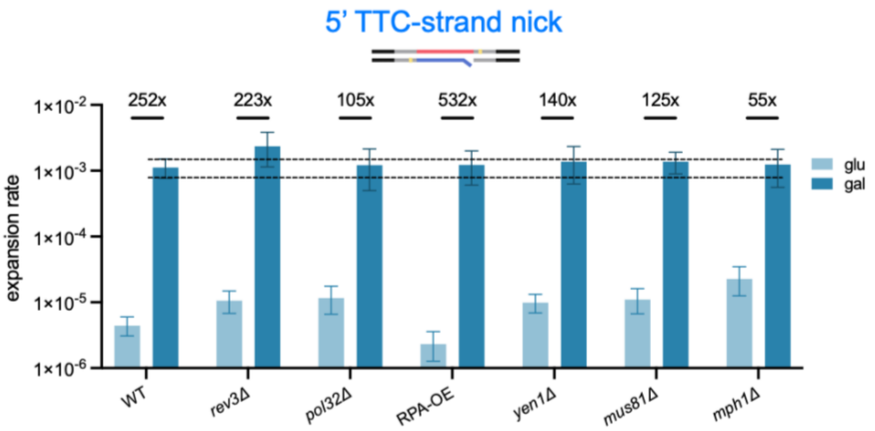

**Figure S3. 5' TTC-strand nick-induced expansion rate of additional genetic controls.** Effect of various mutants on expansion rate of without (glu, light blue) or with (gal, dark blue) galactose induced nCas9-D10A expression. Numbers on top of each genotype show fold change of expansion rate between glucose and galactose group. Error bars represent 95% confidence intervals. Dashed lines represent 95% confidence interval of WT with 5' TTC-strand nick.

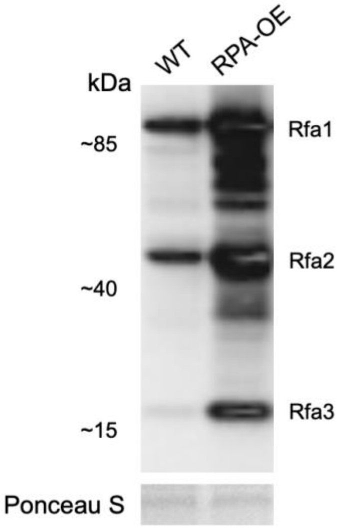

**Figure S4. Western blot showing abundance of all three subunits of yeast RPA in the WT or strain carrying RPA-OE plasmid.** Western blot was probed using anti-RPA antibody. Ladder: BenchMark™ Pre-stained Protein Ladder (Invitrogen #10748010). Ponceau S staining shows protein loading in the same order as Western blot.

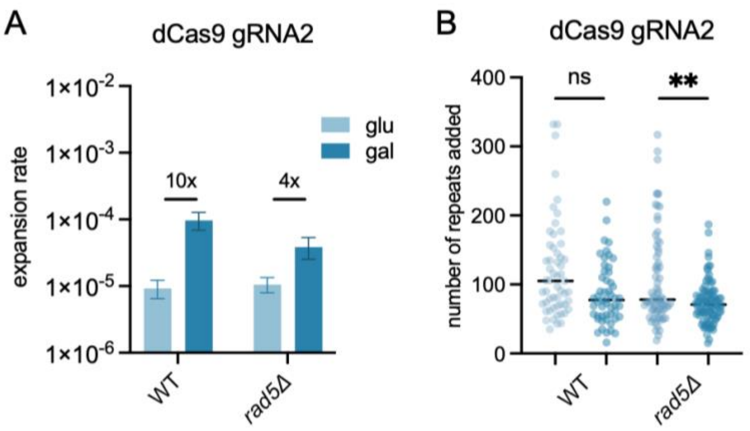

**Figure S5. Effect of deleting *rad5* on dCas9-gRNA2 induced expansion rate and scale.**

(A) Expansion rate of WT or *rad5Δ* strains carrying dCas9 and gRNA2 plasmids without (glu, light blue) or with (gal, dark blue) galactose induction of Cas9 expression. Numbers on top of each genotype show fold change of expansion rate between glucose and galactose group. Error bars represent 95% confidence intervals.

(B) Expansion scale of WT or *rad5Δ* strains carrying dCas9 and gRNA2 plasmids without (glu, light blue) or with (gal, dark blue) galactose induction of Cas9 expression. Dashed lines represent median of distribution. Statistical significance between glucose and galactose groups was calculated via Kolmogorov–Smirnov test.

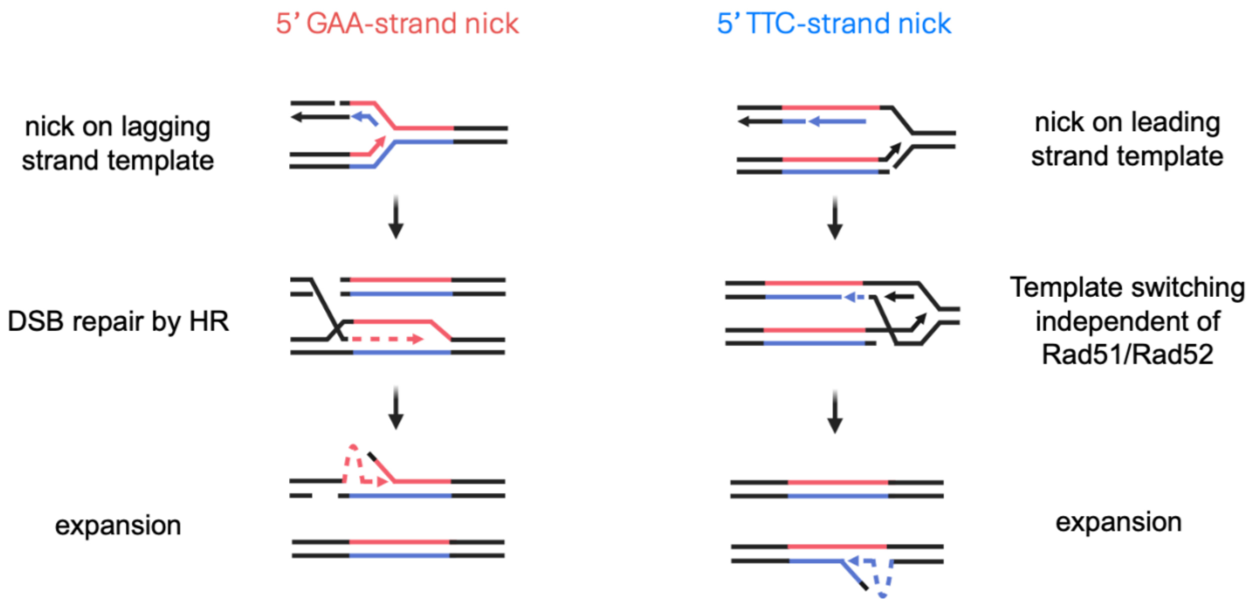

**Figure S6. Alternative model of 5’ nick-mediated $(GAA)_n$ repeat expansion.** Left: The 5’ GAA-strand nick on the lagging strand template right after replication. The nick can be converted into a two-ended DSB before the Okazaki fragment is ligated, which is repaired by HR. Expansion occurs through misalignment of the repeat tract during strand invasion. Right: The 5’ TTC-strand nick on the leading strand template right after replication. The nick is repaired by template switching independent of Rad51/Rad52 using the single-stranded lagging strand template. Expansion could occur via misalignment or flap displacement.

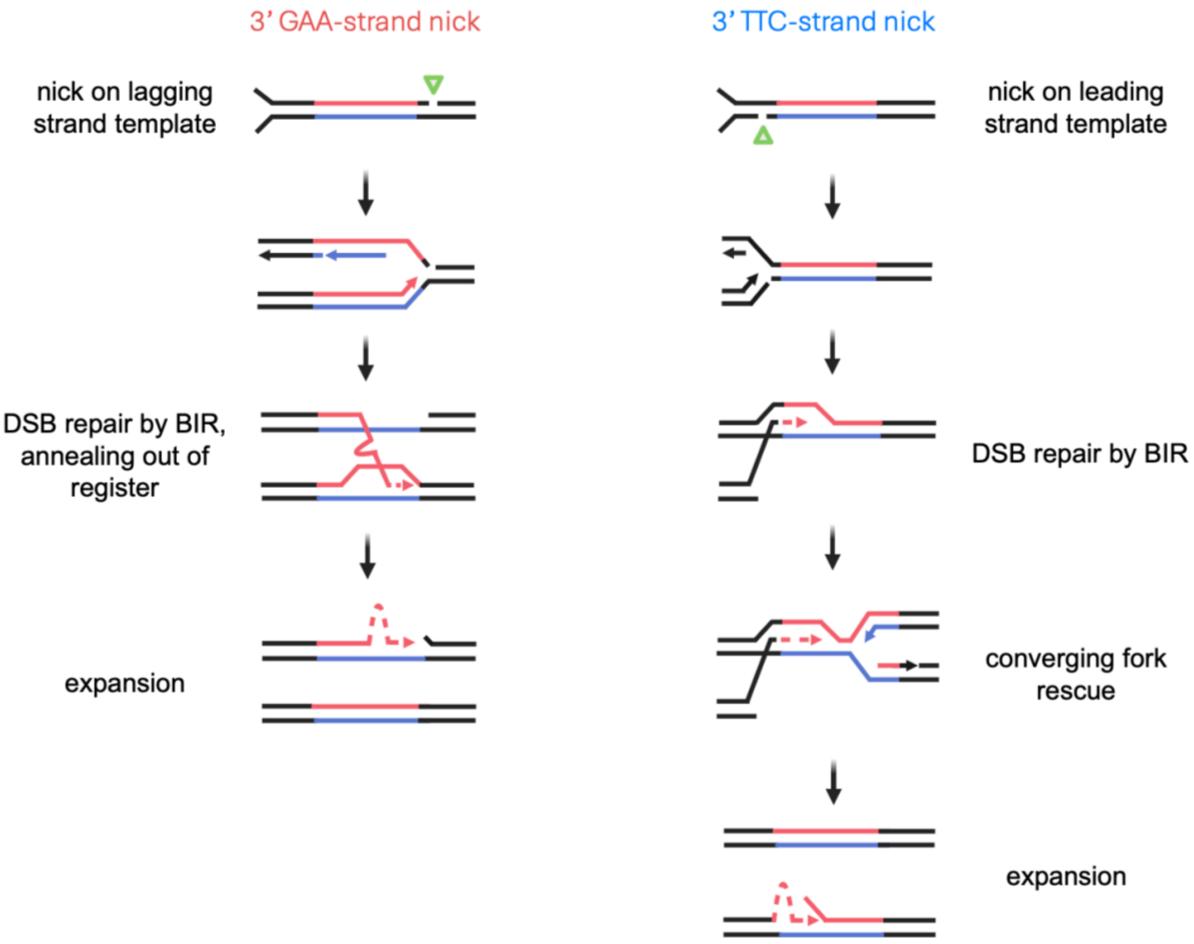

**Figure S7. Model of 3’ nick-mediated $(GAA)_n$ repeat expansion.** Left: 3’ GAA-strand nick on the lagging strand template is converted into a two-ended DSB during replication, which is repaired by BIR. Expansion occurs through misalignment. Right: 3’ TTC-strand nick on the leading strand template is converted into a one-ended DSB during replication when the fork collapses due to CMG helicase falling off. Sister chromatids are held together by cohesin. The one-ended DSB is repaired via BIR and subsequently rescued by a converging fork. Expansion could occur via misalignment or flap displacement.

### Supplementary tables

**Table S1. Primers used in this project**

| **Primer name** | **Primer sequence** | **Notes** |
|---|---|---|
| For construction and checking of the cassettes | | |
| 5'misc-F | GATGGTACGACGGTTTGTAATAGCG | |
| 3'misc-R | AAGCACTAACGATTGCGTGATGG | |
| *TRP1_PPP_R* | CTCTGCAAGCCGCAAACT | |
| *URA3-RT-UnSpl-R* | GAGCCCTTGCATGACAATTC | |
| A36b-F | acgtgtacagttctctttacatcatc | |
| A36a-R | agggtcgttgccttctggtgtt | |
| A36a_p-F | aagactgcaacatactactcagtgc | |
| A36b_p-R | cgacatgatttatcttcgtttcctg | |
| GAA_URA3-F | agaagaattgcacggtcccaa | |
| 3'_UAS-100_2-R | AGGGCTCTCAAGGGCATC | |
| GAA_seq_F | CCCAGGTATTGTTAGCGGtTT | |
| 5_long_F | GATGGTACGACGGTTTGTAATAGCGTTTAAGTAGTTTCCAGTTGG | |
| 3_long_R | AAATAAAAAGCACTAACGATTGCGTGATGGAAGTGGAGAGG | |
| FII_A3_CCCTT84_UR_LR | cttagacccaCGCGGATCCGGAGATCTAGCTTTTCAATTCAATTC | |
| FII_A3_CCCTT84_UR_LF | ttgttatttcCGTCGACGGGTAATAACTGATATAATTAAATTGAAGCTC | |
| FI_pJH21backbone_LF | cggatccgcgTGGGTCTAAGAAACCATTATTATCATG | |
| FI_pJH21backbone_LR | cccgtcgacgGAAATAACAATGTCTGCGAG | |
| JK167_CAN1_fwd | GGCCGCTCTAGAACTAGTGGAT | |
| For amplification of the pRCC-N plasmid with target gRNA sequence | | |
| Rad27_2E1_pRCC_F | TGACAAAGCGGTCTTCAAGAgttttagagctagaaatagcaagttaaaataagg | *rad27-2E1* |
| Rad27_2E1_pRCC_R | TCTTGAAGACCGCTTTGTCAcgatcatttatctttcactgcggag | *rad27-2E1* |
| Chl1_pRCC_F | TCCCTATTACGCCTCGAGGGgttttagagctagaaatagcaagttaaaataagg | *chl1Δ* |
| Chl1_pRCC_R | CCCTCGAGGCGTAATAGGGAcgatcatttatctttcactgcggag | *chl1Δ* |
| rad52_pRCC_F | ACCAGGTTCTTCGTCGAGTCgttttagagctagaaatagcaagttaaaataagg | *rad52Δ* |
| rad52_pRCC_R | GACTCGACGAAGAACCTGGTcgatcatttatctttcactgcggag | *rad52Δ* |

| | | |
|---|---|---|
| Exo1_pRCC_F | TAAGTCAAGCCAAGCTCGGCgttttagagctagaaatagcaagttaaaataagg | *exo1Δ* |
| Exo1_pRCC_R | GCCGAGCTTGGCTTGACTTAcgatcatttatctttcactgcggag | *exo1Δ* |
| Sgs1_pRCC_F | CCCATCTCTTCCAGCACGGCgttttagagctagaaatagcaagttaaaataagg | *sgs1Δ* |
| Sgs1_pRCC_R | GCCGTGCTGGAAGAGATGGGcgatcatttatctttcactgcggag | *sgs1Δ* |
| rad1_pRCC_F | CAACTAAAGTAGGCCCCTGAgttttagagctagaaatagcaagttaaaataagg | *rad1Δ* |
| rad1_pRCC_R | TCAGGGGCCTACTTTAGTTGcgatcatttatctttcactgcggag | *rad1Δ* |
| rad51_pRCC_F | TCAGCGGTTCGGATTAGACCgttttagagctagaaatagcaagttaaaataagg | *rad51Δ* |
| rad51_pRCC_R | GGTCTAATCCGAACCGCTGAcgatcatttatctttcactgcggag | *rad51Δ* |
| Mre11_H125N_pRCC_F | AGGTAATCATGATGATGCGTGTTTTAGAGCTAGAAATAGCAAGTTAAAATAAGG | *mre11Δ* |
| Mre11_H125N_pRCC_R | ACGCATCATCATGATTACCTCGATCATTTATCTTTCACTGCGGAG | *mre11Δ* |
| Rev3_pRCC_F | GATTTGGATATCGCTTACGAgttttagagctagaaatagcaagttaaaataagg | *rev3Δ* |
| Rev3_pRCC_R | TCGTAAGCGATATCCAAATCcgatcatttatctttcactgcggag | *rev3Δ* |
| Rad18_pRCC_F | TAAATACTTATCCACATGTGgttttagagctagaaatagcaagttaaaataagg | *rad18Δ* |
| Rad18_pRCC_R | CACATGTGGATAAGTATTTAcgatcatttatctttcactgcggag | *rad18Δ* |
| Rad54_pRCC_F | ATTATTAAGACAGGGCCCGCgttttagagctagaaatagcaagttaaaataagg | *rad54Δ* |
| Rad54_pRCC_R | GCGGGCCCTGTCTTAATAATcgatcatttatctttcactgcggag | *rad54Δ* |
| Pol32_del_pRCC_F | TTCTTTAGGTGAGTTGGGCTgttttagagctagaaatagcaagttaaaataagg | *pol32Δ* |
| Pol32_del_pRCC_R | AGCCCAACTCACCTAAAGAAcgatcatttatctttcactgcggag | *pol32Δ* |
| Rad5_pRCC_F | TTATTATCTTGCCAAGTGCTgttttagagctagaaatagcaagttaaaataagg | *rad5Δ* |
| Rad5_pRCC_R | AGCACTTGGCAAGATAATAAcgatcatttatctttcactgcggag | *rad5Δ* |
| SRS2_pRCC_F | TTAATCTTCTCACCGGTAGCgttttagagctagaaatagcaagttaaaataagg | *srs2Δ* |
| SRS2_pRCC_R | GCTACCGGTGAGAAGATTAAcgatcatttatctttcactgcggag | *srs2Δ* |
| mph1_pRCC_F | AGAAAAGGGGTTCCGACTCGgttttagagctagaaatagcaagttaaaataagg | *mph1Δ* |

| | | |
|---|---|---|
| mph1_pRCC_R | CGAGTCGGAACCCCTTTTCTcgatcatttatctttcactgcggag | *mph1Δ* |
| Yen1_pRCC_F | CGATTACAATCGTGGAGTCAgttttagagctagaaatagcaagttaaaataagg | *yen1Δ* |
| Yen1_pRCC_R | TGACTCCACGATTGTAATCGcgatcatttatctttcactgcggag | *yen1Δ* |
| Mus81_pRCC_F | AGAGATAAGGCAACTCGCTTgttttagagctagaaatagcaagttaaaataagg | *mus81Δ* |
| Mus81_pRCC_R | AAGCGAGTTGCCTTATCTCTcgatcatttatctttcactgcggag | *mus81Δ* |
| For amplification of a template for repair in CRISPR-Cas9 gene editing | | |
| Rad27_2E1_repairtemp_F | TTTGAAATCTCATGAGTTGACAAAGCGGTCTTCAAGAGAGGTGGAAACAGAAAAAAACTGGCAGAGGCAACAACAGAATTGGAAAAGATGAAGC | *rad27-2E1* |
| Rad27_2E1_repairtemp_R | AATTTTTGGGCTTCTTCATTATGCTCTTTTGAGACTTCCACCAATCTTCTTTCTTGCTTCATCTTTTCCAATTCTGTTGTTGCCTCTGCCAGTTT | *rad27-2E1* |
| Chl1_repairtemp_F | GTTTGTTTCCTTAAAACCCAAAAGAGTAGAAAACCAGGCTAAAAACAGTCACACTAGTCCAAGGAATACGTTTAC | *chl1Δ* |
| Chl1_repairtemp_R | TCAGTTTAGTTTACTATAATATATAGTAGTAATCACAGTATACACGTAAACGTATTCCTTGGACTAGTGTGACTG | *chl1Δ* |
| rad52_repairtemp_F | TTAGTCTGTTAAGAAAAGACGAAAAATATAGCGGCGGGCGGGTTACGCGACCGGTATCGAAACGCTTCCTGGCCG | *rad52Δ* |
| rad52_repairtemp_R | AGGATTTTGGAGTAATAAATAATGATGCAAATTTTTTATTTGTTTCGGCCAGGAAGCGTTTCGATACCGGTCGCG | *rad52Δ* |
| Exo1_repairtemplate | ACCACATTAAAATAAAAGGAGCTCGAAAAAACTGAAAGGCGTAGAAAGGAAAGTTAAGTACTGCACGTTTCATATCGGAGGTATATTTTTCAAATGAAAA | *exo1Δ* |
| Sgs1_repairtemplate | ATTATTGTTGTATATATTTAAAAAATCATACACGTACACACAAGGCGGTAAGAGTAGAAAAATAAATAGTGTTACTTATAACTACGACACCATTCGCCAA | *sgs1Δ* |
| rad1_del_repairtemp_F | TAAATGTGTAAAAATAATATTGCACTATCCTGTTGAAAATATCTTTCCAGAATATTGTTTAATTTAACGA | *rad1Δ* |
| rad1_del_repairtemp_R | TCGCATTTTATACTGATGTTTTAACAGGGTTCGTTAAATTAAACAATATTCTGGAAAGATATTTTCAACA | *rad1Δ* |
| rad51_repairtemp_F | CGACAAAGAGCAGACGTAGTTATTTGTTAAAGGCCTACTAATTTGTTATCGTCATGTATTTGGTCTCTTG | *rad51Δ* |

| | | |
|---|---|---|
| rad51_repairtemp_R | AGAGGAGAATTGAAAGTAAACCTGTGTAAATA AATAGAGACAAGAGACCAAATACATGACGATA ACAAAT | *rad51Δ* |
| mre11_del_repairtemp_F | GCAGACAATTGACGCAAGTTGTACCTGCTCAG ATCCGATAAAACTCGACTTTGTACTTGATCCCTA TATT | *mre11Δ* |
| mre11_del_repairtemp_R | AAGCCCTTGGTTATAAATAGGATATAATATAATAT AGGGATCAAGTACAAAGTCGAGTTTTATCGGAT CT | *mre11Δ* |
| Rev3_repairtemp_F | AAAGTATTTGAGTCAATACAAAACTACAAGTTG TGGCGAAATAAAATGTTTGGAATCTAGACACAG ATAT | *rev3Δ* |
| Rev3_repairtemp_R | TATACATAGAAACAAATAACTACTCATCATTTTG CGAGACATATCTGTGTCTAGATTCCAAACATTTT AT | *rev3Δ* |
| Rad18_repairtemp_F | AAACCATCCGCAAGTGAGCATCACAGCTACTAA GAAAAGGCCATTTTTACTACTCGGTGTGTATATG TAA | *rad18Δ* |
| Rad18_repairtemp_R | ATTAATTAACAAATGTGCACAAGCTAACAAACA GGCCTGATTACATATACACACCGAGTAGTAAAA ATGG | *rad18Δ* |
| Rad54_repairtemp_F | AAACGCTCAGAACTTAGCTCTATTTCAAGGTAC CATATATATTTCCTTATAACTGTCTCTTACATACA TG | *rad54Δ* |
| Rad54_repairtemp_R | CGACGATCGAATTCTACTTTTTGTTTTTGTTTTA TAAGTACATGTATGTAAGAGACAGTTATAAGGA AAT | *rad54Δ* |
| Pol32_del_repairtemp_F | TCAGCTCGAAATAATATTTCACATTAACTAACAA CCAGAAATAGGCTTTAGTTAACTCAATCGGTAA TTACCAAACACTTTCCATTACTAATTGT | *pol32Δ* |
| Pol32_del_repairtemp_R | TATTTTTCTATCACGTAAGTTGACATTTGTATTAT ACATTACATCACAATTAGTAATGGAAAGTGTTT GG TAATTACCGATTGAGTTAACTAAAG | *pol32Δ* |
| Rad5_repairtemp_F | AGTTACATTATCAAAAGGCCTTAGAAACACACC TAAAGTCTTACAGTATCACAATCAGACAAACAG CGTC | *rad5Δ* |
| Rad5_repairtemp_R | TTTCTTCTATGCTATCTTGTATGATAAATCTCATA ACTTTGACGCTGTTTGTCTGATTGTGATACTGTA A | *rad5Δ* |
| srs2_repairtemp_F | GAGTATCATTCCAATTTGATCTTTCTTCTACCGG TACTTAGGGATAGCAATAGCACTTTCATGCCTG ACT | *srs2Δ* |
| srs2_repairtemp_R | AAATTATAAACCGCCTCCAATAGTTGACGTAGT CAGGCATGAAAGTGCTATTGCTATCCCTAAGTA CCGG | *srs2Δ* |

| | | |
|---|---|---|
| mph1_repairtemp_F | ATTCAACACATTCCGGTTCTGTTTTATTTTAGTGTCCTTTTTTCTCTCTGTAGGAGACTCTTATACGTCG | *mph1Δ* |
| mph1_repairtemp_R | ATTACAGCAGCGTTATTTTGTATAGACGCCGACGTATAAGAGTCTCCTACAGAGAGAAAAAAGGACACT | *mph1Δ* |
| Yen1_repairtemp_F | ATGACAGTTCTATTGCATTTTACCTACTTGTATATTCTGGATACTGCACAAGAAAGTTAACGGGCACAGC | *yen1Δ* |
| Yen1_repairtemp_R | CGTTTTCGGCGCGATCAACTGTGGTGGCGGATTTTTTGACGCTGTGCCCGTTAACTTTCTTGTGCAGTAT | *yen1Δ* |
| Mus81_repairtemp_F | GGCGTAAACAAAGTTTCAAAGGATTGATACGAACACACATTCCTAGCATGAAAGCCAATTTGAAATATAG | *mus81Δ* |
| Mus81_repairtemp_R | ATATCATCACTTTTTTCTTTATAAAACCTTGCAGGGATGACTATATTTCAAATTGGCTTTCATGCTAGGA | *mus81Δ* |
| For Sanger sequencing of the missense mutations | | |
| Rad27_2E1_trans_F | GCCTTGTTATGTCTTCGACGG | *rad27-2E1* |
| Rad27_2E1_trans_R | AGCACATTGAGCCTCAGCTT | *rad27-2E1* |
| For knockout fragments amplification | | |
| Rad10KOpAGfwd | ATGAACAATACTGATCCTACTTCATTTGAAAGTATATTGGCTGGTGTGGCCAGCTGAAGCTTCGTACGC | *rad10::NAT* |
| Rad10KOpAGrev | TAAATTCAAATATTCAATATATTTTGCAGCTTGTTCAAAATTAAATGCCATAGGCCACTAGTGGATCTG | *rad10::NAT* |
| For checking knockout (internal) | | |
| CHL1-F | CCCTCGTTCCCATCATCCTT | *chl1Δ* |
| CHL1-R | CGTATCAGGACAATGACGCC | *chl1Δ* |
| RAD52_chk_in_F | GCCAAGAAATCTGCCGTTAC | *rad52Δ* |
| RAD52_chk_in_R | TGAGCTTTCGCTGATTTCATCC | *rad52Δ* |
| Rad51-chk_in_F | agatcggagctgatttgtttgac | *rad51Δ* |
| Rad51-chk_in_R | cttcaccgccaccaatatcc | *rad51Δ* |
| Exo1-F | CAGCGGGAGGGAAAACTGAT | *exo1Δ* |
| Exo1-R | CTCTGTTGGCTAGAGGTTGGTG | *exo1Δ* |
| JK341_Sgs1_int_fwd | tgcaaactttgtcgaacgatac | *sgs1Δ* |
| JK342_Sgs1_int_rev | cgacaagagaactagccatgtg | *sgs1Δ* |
| RAD1-F | CGAACTGGCACCGAATTTCT | *rad1Δ* |
| RAD1-R | CTAAAGTAGGCCCCTGAAGG | *rad1Δ* |

| | | |
|---|---|---|
| JK254_Mre11_int_fwd | tggatatacttcatgcgactgg | *mre11Δ* |
| JK255_Mre11_int_rev | atgaccccatatcaccatatcc | *mre11Δ* |
| REV3-F | GCATGCACACCCCTCATAGTAAGT | *rev3Δ* |
| Rev3 2400B | tggcatttgactctggcaagttcc | *rev3Δ* |
| RAD18-F | CCACTGAGTTCCAAACCATC | *rad18Δ* |
| RAD18-R | GACTTCTGGAGTTCGTACCT | *rad18Δ* |
| Rad54_in_F | GTCCCTGTGGTTATTGATCC | *rad54Δ* |
| Rad54_in_R | TACACTGCAATGTCTTACCC | *rad54Δ* |
| pol32_in_F | gaccacgccagaagaaacaa | *pol32Δ* |
| pol32_in_R | gctgtcgtttccaacaagtc | *pol32Δ* |
| Rad5_in_F | ctgaagtccataacaatctccga | *rad5Δ* |
| Rad5_in_R | gccatttgaactgcttccataa | *rad5Δ* |
| JK300_SRS2_int_fwd | TCTAGTCAGCAAACGAAAGGTG | *srs2Δ* |
| JK301_SRS2_int_rev | AGGCAAACTTTATTATAC | *srs2Δ* |
| MPH1_chk_in_F | GGTCATGGGAAGTTACAATGTGTT | *mph1Δ* |
| MPH1_chk_in_R | CGTCAGCTACCGAGTCTATGAA | *mph1Δ* |
| Yen1_F | cCACAAGACTGACCACCGTCG | *yen1Δ* |
| Yen1_R | cCACAAGACTGACCACCGTCG | *yen1Δ* |
| JK260_MUS81_int_fwd | CACAGCAAATCTGACTGACCTC | *mus81Δ* |
| JK261_MUS81_int_rev | TTCGAAATCACCACTACACCAC | *mus81Δ* |
| For checking knockout (external) | | |
| CHL-upstr-F | GGCACTACTGCAACTTCAGT | *chl1Δ* |
| CHL1-dnstr-R2 | CTGAAACTGTAACATCCACC | *chl1Δ* |
| Rad52_up_F | CGGTGAGTGTGGCAACGCC | *rad52Δ* |
| Rad52_down_R | CAATGAACCTAAGGATTCC | *rad52Δ* |
| Rad51_up_F | CTAGGCCACACTTCGTTACC | *rad51Δ* |
| Rad51_down_R | CCAATTTCAGGGTATGCACC | *rad51Δ* |
| exo1_KOchk_Fwd | GTATTACGTCCAAACTAAGTTCGCG | *exo1Δ* |
| exo1_KOchk_Rev | GACCGCTAGCGGCTTGATTAG | *exo1Δ* |
| JK125_SGS1_fwd | TATCATCCTCATCATCCAG | *sgs1Δ* |
| JK126_SGS1_rev | CTCGAGCCTGATCTAAAAGCTG | *sgs1Δ* |
| RAD1_Chk_F | ATGTAATCAACCTGTCCCGTCC | *rad1Δ* |
| RAD1_Chk_R | AGTGGAAGATGAATTGCGGATGA | *rad1Δ* |
| JK232_Mre11_fwd | ccaatcatttcgaccgtcactc | *mre11Δ* |
| JK233_Mre11_rev | cacaaggggacggttaatgagg | *mre11Δ* |
| REV3-upstr-F | CGAGTGCAGTGCGTCTAGAAATAGTGT | *rev3Δ* |
| Rev3_down_R | GGCGTTATTAATGCATCTGGGTCC | *rev3Δ* |
| RAD18-upstr-F | GAGCAATGCCACATTAGAAG | *rad18Δ* |
| Rad18_down_R | TATTTCAGCACTTAACGTGG | *rad18Δ* |

| | | |
|---|---|---|
| Rad54_up_F | TCTCACTTGACGTAATAGCC | *rad54Δ* |
| Rad54_down_R | CTTTGGCAAGAATTTCATCC | *rad54Δ* |
| JK225_pol32_fwd | tttccactacggtgtaactttcc | *pol32Δ* |
| JK226_POL32_rev | TGTCCTTCGGATGGTATATTAGG | *pol32Δ* |
| RAD5del-chk-F | TTACGCGTCATAAACCCTT | *rad5Δ* |
| RAD5del-chk-R | ACAGCATCTGGATTTCTTCA | *rad5Δ* |
| srs2_up_F | ATACTACTGCTTAGGCTACC | *srs2Δ* |
| srs2_down_R | GTCACGATAGATTCTTCACC | *srs2Δ* |
| mph1_ up_F | TGTTGGTGGCTCGCTATTAG | *mph1Δ* |
| mph1_down_R | CAGATTGTACTCGTCGTTGG | *mph1Δ* |
| Yen1_upstr_F | TATTGGCATTGAACACTGGC | *yen1Δ* |
| Yen1_dnstr_R | GGTTTTTGAAATAAGCGACGAC | *yen1Δ* |
| JK248_MUS81_fwd | agaggtggtggtcaaatcatcc | *mus81Δ* |
| JK249_MUS81_rev | actgcctccaattttgattgcc | *mus81Δ* |
| for checking repeat length | | |
| A2 | CTCGATGTGCAGAACCTGAAGCTTGATCT | |
| B2 | GCTCGAGTGCAGACCTCAAATTCGATGA | |
| UC1 | ggtcccaattctgcagatatccatcacac | |
| UC6 | GCAAGGAATGGTGCATGCTCGAT | |
| for construction of Cas9 and gRNA plasmids | | |
| gRNA_seq | TTTTGTAGTGCCCTCTTGGGC | |
| Ori_F | tgtgatgctcgtcaggggggg | |
| GAA_gRNA1_F | CTTCTTCTTCATCGAATTTGGTTTT | gRNA1 |
| GAA_gRNA1_R | CAAATTCGATGAAGAAGAAGGATCA | gRNA1 |
| GAA_gRNA2_F | GAAGAAGAAGATCAAGCTTCGTTTT | gRNA2 |
| GAA_gRNA2_R | GAAGCTTGATCTTCTTCTTCGATCA | gRNA2 |
| NcoI_Cas9_F | tgtctccATGGACAAGAAGTACTCCATTG | |
| Cas9_R | CGTAATTGACTGATGAATCAGTGTG | |

**Table S2. Yeast strains used in this study**

| **Strain** | **Genotype** | **notes** |
|---|---|---|
| CH1585 | *MAT**a**, leu2Δ1, trp1Δ63, ura3Δ52, his3Δ200* | Shishkin et al. 2009 |
| SMY706 | CH1585 *MATa, leu2Δ1, trp1Δ63, ura3Δ52, his3Δ200* MIP1, HAP1 | |
| SMY923 | SMY706 *ura3Δ* | |
| LLY70/71 | SMY923 *ChrIII(75594-75641)::UR-(GAA)100-A3-TRP1* | |

| | |
|---|---|
| LLY73/74 | LLY70 pRS415_pGal_nCas9(D10A) pSK26_GAA_gRNA1_HYG |
| LLY75/76 | LLY70 pRS415_pGal_nCas9(D10A) pSK26_GAA_gRNA2_HYG |
| LLY160/161 | LLY70 pRS415_pGal_nCas9(H840A) pSK26_GAA_gRNA1_HYG |
| LLY186/187 | LLY70 pRS415_pGal_nCas9(H840A) pSK26_GAA_gRNA2_HYG |
| LLY78 | LLY70 *ChrIII(75594-75641)::UR-(GAA)28-A3-TRP1* |
| LLY80 | LLY70 *ChrIII(75594-75641)::UR-(GAA)40-A3-TRP1* |
| LLY81 | LLY70 *ChrIII(75594-75641)::UR-(GAA)33-A3-TRP1* |
| LLY82/83 | LLY78 pRS415_pGal_nCas9(D10A) pSK26_GAA_gRNA1_HYG |
| LLY84/85 | LLY81 pRS415_pGal_nCas9(D10A) pSK26_GAA_gRNA1_HYG |
| LLY105/106 | LLY80 pRS415_pGal_nCas9(D10A) pSK26_GAA_gRNA1_HYG |
| LLY108/109 | LLY80 pRS415_pGal_nCas9(D10A) pSK26_GAA_gRNA2_HYG |
| LLY98/LLY99 | LLY70 *rad27-2E1* |
| LLY115/116 | LLY98 pRS415_pGal_nCas9(D10A) pSK26_GAA_gRNA1_HYG |
| LLY123/124 | LLY70 pUC57-2u-HIS3-RFA1-RFA2-RFA3 |
| LLY125/126 | LLY123 pRS415_pGal_nCas9(D10A) pSK26_GAA_gRNA1_HYG |
| LLY170/171 | LLY123 pRS415_pGal_nCas9(D10A) pSK26_GAA_gRNA2_HYG |
| LLY129/130 | LLY70 *chl1Δ* |
| LLY133/134 | LLY70 *pol32Δ* |
| LLY142/143 | LLY129 pRS415_pGal_nCas9(D10A) pSK26_GAA_gRNA1_HYG |
| LLY145/146 | LLY129 pRS415_pGal_nCas9(D10A) pSK26_GAA_gRNA2_HYG |
| LLY148/149 | LLY133 pRS415_pGal_nCas9(D10A) pSK26_GAA_gRNA1_HYG |
| LLY151/152 | LLY133 pRS415_pGal_nCas9(D10A) pSK26_GAA_gRNA2_HYG |
| LLY154/155 | LLY135 pRS415_pGal_nCas9(D10A) pSK26_GAA_gRNA1_HYG |
| LLY157/158 | LLY135 pRS415_pGal_nCas9(D10A) pSK26_GAA_gRNA2_HYG |
| LLY163/164 | LLY70 pJA29_pGal_dCas9 +pSK26_GAA_gRNA1_HYG |
| LLY166/167 | LLY70 pJA29_pGal_dCas9 +pSK26_GAA_gRNA2_HYG |

| | |
|---|---|
| LLY176 | LLY70 *rad52Δ* |
| LLY209/210 | LLY176 pRS415_pGal_nCas9(D10A) pSK26_GAA_gRNA1_HYG |
| LLY222/223 | LLY176 pRS415_pGal_nCas9(D10A) pSK26_GAA_gRNA2_HYG |
| LLY216 | LLY70 *rad51Δ* |
| LLY224/225 | LLY216 pRS415_pGal_nCas9(D10A) pSK26_GAA_gRNA1_HYG |
| LLY227/228 | LLY216 pRS415_pGal_nCas9(D10A) pSK26_GAA_gRNA2_HYG |
| LLY248 | LLY70 *rad1Δ* |
| LLY252 | LLY70 *mre11Δ* |
| LLY254/255 | LLY176 pRS415_pGal_nCas9(H840A) pSK26_GAA_gRNA1_HYG |
| LLY257/258 | LLY176 pRS415_pGal_nCas9(H840A) pSK26_GAA_gRNA2_HYG |
| LLY260/261 | LLY216 pRS415_pGal_nCas9(H840A) pSK26_GAA_gRNA1_HYG |
| LLY263/264 | LLY216 pRS415_pGal_nCas9(H840A) pSK26_GAA_gRNA2_HYG |
| LLY277/278 | LLY252 pRS415_pGal_nCas9(D10A) pSK26_GAA_gRNA1_HYG |
| LLY280/281 | LLY252 pRS415_pGal_nCas9(D10A) pSK26_GAA_gRNA2_HYG |
| LLY286 | LLY70 *rad10::NAT* |
| LLY289 | LLY248 *rad10::NAT* |
| LLY291 | LLY70 *srs2Δ* |
| LLY294/295 | LLY248 pRS415_pGal_nCas9(D10A) pSK26_GAA_gRNA1_HYG |
| LLY297/298 | LLY286 pRS415_pGal_nCas9(D10A) pSK26_GAA_gRNA1_HYG |
| LLY300/301 | LLY289 pRS415_pGal_nCas9(D10A) pSK26_GAA_gRNA1_HYG |
| LLY311/312/313 | LLY291 pRS415_pGal_nCas9(D10A) pSK26_GAA_gRNA1_HYG |
| LLY314/315 | LLY291 pRS415_pGal_nCas9(D10A) pSK26_GAA_gRNA2_HYG |
| LLY317/318 | LLY248 pRS415_pGal_nCas9(D10A) pSK26_GAA_gRNA2_HYG |
| LLY320/321 | LLY286 pRS415_pGal_nCas9(D10A) pSK26_GAA_gRNA2_HYG |
| LLY323/324 | LLY289 pRS415_pGal_nCas9(D10A) pSK26_GAA_gRNA2_HYG |
| LLY336 | LLY70 *yen1Δ* |

| | |
|---|---|
| LLY339 | LLY70 *mus81Δ* |
| LLY342 | LLY70 *rad5Δ* |
| LLY347/348 | LLY133 pRS415_pGal_nCas9(H840A) pSK26_GAA_gRNA1_HYG |
| LLY349/350 | LLY133 pRS415_pGal_nCas9(H840A) pSK26_GAA_gRNA2_HYG |
| LLY351/352 | LLY336 pRS415_pGal_nCas9(D10A) pSK26_GAA_gRNA1_HYG |
| LLY353/354 | LLY336 pRS415_pGal_nCas9(D10A) pSK26_GAA_gRNA2_HYG |
| LLY356 | LLY339 pRS415_pGal_nCas9(D10A) pSK26_GAA_gRNA1_HYG |
| LLY357/358 | LLY339 pRS415_pGal_nCas9(D10A) pSK26_GAA_gRNA2_HYG |
| LLY359/360 | LLY342 pRS415_pGal_nCas9(D10A) pSK26_GAA_gRNA1_HYG |
| LLY361/362 | LLY342 pRS415_pGal_nCas9(D10A) pSK26_GAA_gRNA2_HYG |
| LLY375 | LLY70 *mph1Δ* |
| LLY384 | LLY336 *mus81Δ* |
| LLY385/386 | LLY384 pRS415_pGal_nCas9(D10A) pSK26_GAA_gRNA1_HYG |
| LLY388/389 | LLY384 pRS415_pGal_nCas9(D10A) pSK26_GAA_gRNA2_HYG |
| LLY419 | LLY70 *exo1Δ* |
| LLY422/423 | LLY135 pRS415_pGal_nCas9(H840A) pSK26_GAA_gRNA1_HYG |
| LLY424/425 | LLY135 pRS415_pGal_nCas9(H840A) pSK26_GAA_gRNA2_HYG |
| LLY431/432 | LLY419 pRS415_pGal_nCas9(D10A) pSK26_GAA_gRNA1_HYG |
| LLY435 | LLY70 *sgs1Δ* |
| LLY438 | LLY384 *exo1Δ* |
| LLY441 | LLY70 *rev3Δ* |
| LLY444 | LLY70 *rad18Δ* |
| LLY447/448/449 | LLY435 pRS415_pGal_nCas9(D10A) pSK26_GAA_gRNA1_HYG |
| LLY468/469 | LLY435 pRS415_pGal_nCas9(D10A) pSK26_GAA_gRNA2_HYG |
| LLY451/452 | LLY438 pRS415_pGal_nCas9(D10A) pSK26_GAA_gRNA2_HYG |
| LLY453 | LLY70 *rad54Δ* |

| | |
|---|---|
| LLY457/458 | LLY444 pRS415_pGal_nCas9(D10A) pSK26_GAA_gRNA1_HYG |
| LLY460/461 | LLY444 pRS415_pGal_nCas9(D10A) pSK26_GAA_gRNA2_HYG |
| LLY463/464 | LLY453 pRS415_pGal_nCas9(D10A) pSK26_GAA_gRNA1_HYG |
| LLY466/467 | LLY453 pRS415_pGal_nCas9(D10A) pSK26_GAA_gRNA2_HYG |
| LLY470/471 | LLY453 pRS415_pGal_nCas9(H840A) pSK26_GAA_gRNA1_HYG |
| LLY473/474 | LLY453 pRS415_pGal_nCas9(H840A) pSK26_GAA_gRNA2_HYG |
| LLY478/479 | LLY441 pRS415_pGal_nCas9(D10A) pSK26_GAA_gRNA1_HYG |
| LLY482/483 | LLY441 pRS415_pGal_nCas9(D10A) pSK26_GAA_gRNA2_HYG |
| LLY485 | LLY435 *exo1Δ* |
| LLY503/504 | LLY485 pRS415_pGal_nCas9(D10A) pSK26_GAA_gRNA1_HYG |
| LLY505/506 | LLY485 pRS415_pGal_nCas9(D10A) pSK26_GAA_gRNA2_HYG |
| LLY517 | SMY923 *ChrIII(75594-75641)::A3-(TTC)95-UR-TRP1* |
| LLY520 | LLY517 *rad52Δ* |
| LLY523/524 | LLY517 pRS415_pGal_nCas9(D10A) pSK26_GAA_gRNA1_HYG |
| LLY526/527 | LLY517 pRS415_pGal_nCas9(D10A) pSK26_GAA_gRNA2_HYG |
| LLY555/556 | LLY342 pJA29_pGal_dCas9 +pSK26_GAA_gRNA1_HYG |
| LLY558/559 | LLY342 pJA29_pGal_dCas9 +pSK26_GAA_gRNA2_HYG |

**Table S3. Plasmids used in this study**

| Name | Note |
|---|---|
| pRS415_pGal-nCas9(D10A) | from addgene |
| pRS415_pGal-nCas9(H840A) | from addgene |
| pJA29 pGal Cas9dead | DH10b |
| pSK26_3_gRNA_HYG (bRA98-noCas9) | |
| pLZ1_GAA_gRNA1_HYG | DH5a |
| pLZ2_GAA_gRNA2_HYG | DH5a |
| pUC57-2u-HIS3-RFA1-RFA2-RFA3 | from GeneUniversal |
| pYES3-pURA-UR-(GAA)100-A3-TRP1 | |
| pSS1-A3-(TTC)95-UR-pURA-TRP1 | |

**Table S4. Media recipe**

| | | |
|---|---|---|
| Media for maintaining nickase plasmids | | |
| **Hygromycin Dropout media Proline rich** | | |
| YNB w/o AS | 1.7 | g |
| Destrose/Galactose | 20 | g |
| Agar | 25 | g |
| Dropout mix | 2 | g |
| Proline | 1.085 | g |
| H2O | 1000 | ml |
| | autoclave | |
| HygB (50 mg/ml) | 6000 | ul, (300 ug/ml total) |
| Media for starving and maintaining quiescent yeast cells with nickase plasmids | | |
| **Phosphate limited media** | | |
| YNB - AA-AS-Phos-Sugar | 0.71g/L | |
| Potassium Chloride | 1g/L | |
| Proline | 1.085g/L | |
| Dextrose | 20g/L | |
| Dropout -leu | 2g/L | |
| Potassium phosphate | 0.05g/L | |
| HygB | 6mL/L | |
| **No phosphate media** | | |
| Same as above except: | | |
| Dextrose/Galactose | 2g/L | |
| Potassium phosphate | 0g | |
